# Efficacy and safety evaluation of artificial intelligence-identified antimicrobial peptides for use against avian pathogenic *Escherichia coli* in the poultry industry

**DOI:** 10.1101/2025.09.15.675954

**Authors:** Emre Demirsoy, Teagan I. Parkin, Shaeleen E. Mihalynuk, Anna H. Dema, Lorissa Corrie, Marika E. Heilker, Haley N. Kuecks-Winger, Anat Yanai, Uluc B. Birol, Michael McIlwee, Kay de Wet, Mathijs Knipscheer, Victoria Bowes, Vanessa Tuytel, Liam Ritchie, Wolfgang Köster, Emil Berberov, William R. Cox, Monica Kotkoff, Vanessa C. Thompson, Rene L. Warren, Erin Fraser, Linda M.N. Hoang, Fraser Hof, Fatih Birol, Caren C. Helbing, Inanc Birol

**Author notes:** These authors contributed equally to this work (Emre Demirsoy, Teagan I. Parkin).

## Abstract

The overuse of antibiotics in both veterinary and human medicine has resulted in the emergence of antibiotic-resistant bacteria, prompting a search for effective alternatives. Antimicrobial peptides (AMP) are short, often cationic, peptide-based molecules with antimicrobial and immunomodulatory activity, which makes them promising alternatives to conventional antibiotics in poultry production. From a prior machine-learning-guided screen of 875 candidate AMPs, 62 exhibited activity against avian pathogenic *Escherichia coli* (APEC) and low in vitro hemolytic and cytotoxic activity. We selected three lead AMPs from this list (named TeRu4, TeBi1, and PeNi4), and evaluated their in vitro and in vivo efficacy, safety, and immunomodulatory potential for use in poultry farming. In animal experiments, AMPs were administered via in ovo injection on day 18 of embryonic development. In APEC challenge trials, yolk sacs were inoculated with APEC post-hatch to assess early chick mortality, while in pen trials, birds were raised in a commercial production setting for 35 days. For challenged birds, TeBi1 (10 μg/egg) significantly reduced bacterial detection in the air sac and pericardium, increased body weight by 50% and reduced cytokine transcript levels by 10-30% on day 7 post hatch. In HD11 chicken macrophage-like cultured cells, TeRu4 (16 μg/mL) suppressed lipopolysaccharide (LPS)-induced pro-inflammatory cytokine transcript levels. In pen trials, TeRu4 (20 μg/egg) increased the survival probability of female birds by 4.9%, while TeBi1 (20 μg/egg) increased the survival probability of all birds by 4.4%, by day 35. Gene expression analysis revealed AMP- and sex-specific cytokine responses. In pen trials, no significant differences were observed in mean weights, feed conversion ratio (FCR), and flock uniformity on day 35. These findings demonstrate that the three selected AMPs are safe antibiotic alternatives that improve survival, modulate immune responses, and maintain normal growth performance in broiler chickens.

## INTRODUCTION

Avian pathogenic *Escherichia coli* (**APEC**) is a Gram-negative bacterium that poses a severe threat to the poultry industry. APEC infections induce fatal bacterial septicemia and early chick mortality (**ECM**), a significant cause of mortality and morbidity across multiple poultry commodities (Cuperus et al., 2016; Allan et al., 2018; Rezaee et al., 2021; Nguyen et al., 2021; Sarfraz et al., 2022; Hu et al., 2022). Antibiotics have been used to treat bacterial infections since the early 20th century. In the poultry industry, antibiotics have been used as growth promoters and prophylactic agents since the 1950s (Bean-Hodgins and Kiarie, 2021). However, unnecessary or overuse of antibiotics has contributed to the rise of antibiotic-resistant bacteria in both veterinary and human medicine (Bean-Hodgins and Kiarie, 2021; Richter et al., 2022). To address this growing challenge, regulatory, veterinary, and poultry industry policy changes in Canada and in other jurisdictions have been instituted to limit the use of antimicrobials for growth promotion purposes and, also, to limit the use of medically important antimicrobials (Bean-Hodgins and Kiarie, 2021; Wallinga et al., 2022; Schmerold et al., 2023). This has resulted in fewer antimicrobial options being available to control pathogens in poultry flocks (Bean-Hodgins and Kiarie, 2021). The One Health approach recognizes the linkages between human, animal, and environmental health, and highlights the urgency of addressing antibiotic resistance as a global crisis (Destoumieux-Garzón et al., 2018). Unfortunately, novel antibiotics are not approved at a pace sufficient to combat developing antibiotic-resistant bacteria (Richter et al., 2022). Moreover, the last new class of antibiotics, oxazolidinones, was discovered with linezolid in 1987 (Durand et al., 2019). Consequently, there is growing interest in alternative antimicrobial strategies, such as antimicrobial peptides (**AMP**), to address this therapeutic gap.

AMPs are peptide-based biomolecules composed of 5 to 100 amino acids and are typically cationic (Bahar and Ren, 2013). In nature, they are produced by the innate immune system and act as the first line of defense against infections in a wide variety of organisms, including humans, animals, and plants (Bahar and Ren, 2013; Durand et al., 2019). Most conventional antibiotics have specific cellular targets, including cellular machinery involved in DNA and protein synthesis and components of bacterial cell walls (Bahar and Ren, 2013). In contrast, AMPs primarily cause rapid cell death by binding to anionic bacterial cell membranes and disrupting their integrity, leading to cell lysis (Bahar and Ren, 2013; Durand et al., 2019). AMPs typically do not target eukaryotic cells, as their membranes usually have low anionic charge (Bahar and Ren, 2013). However, at higher concentrations, some AMPs may exhibit cytotoxic effects on eukaryotic cells (Bahar and Ren, 2013). Therefore, rigorous safety evaluation of AMPs is essential before regulatory approval, including in vitro assays, such as hemolysis and cytotoxicity tests, as well as in vivo experiments.

AMPs exert less selective pressure on bacteria than conventional antibiotics due to their nonspecific targets and diverse mechanisms of action, thus reducing the development of antimicrobial resistance (Zasloff, 2002; Hancock and Sahl, 2006; Fjell et al., 2012; Richter et al., 2022). In addition to their direct antimicrobial activity, some AMPs demonstrate immunomodulatory properties, recruiting immune cells or regulating cytokine levels to enhance the host response to infection (Rima et al., 2021; Richter et al., 2022). Cytokines are important signaling molecules that facilitate the growth, differentiation, activation of immune cells (Garcia et al., 2021). In the present study, the effect of AMPs on the expression of the following cytokines was investigated: Interleukin (**IL**)-1β, IL-8, IL-6, IL-10, and interferon gamma (**IFN-*γ***), as well as the transcription factor interferon regulator factor 4 (**IRF-4**). IL-1β, IL-8, and IL-6 are associated with activation of inflammation in chickens (Kogut, 2002; Garcia et al., 2021). Furthermore, IL-8 recruits heterophils to the infection site (Kogut, 2002). Conversely, IL-10 plays a role in tissue repair, angiogenesis, and attenuation of the inflammatory response and is an inducible feedback regulator of the immune response in chickens (Rothwell et al., 2004; Wu et al., 2016; Garcia et al., 2021). The roles of IRF-4 and IFN-*γ* are not well-characterized in chickens; however, there is evidence that IRF-4 plays a critical role in B cell development and maturation, similar to its mammalian counterpart (Yu et al., 2021), while IFN-*γ* generally functions to bridge the adaptive and innate immune systems (Yuk et al., 2016; Santhakumar et al., 2017).

Previously, we screened 875 AMP candidates, identified using our artificial intelligence (**AI**) based prediction tools: AMPlify (Li et al., 2022), rAMPage (Lin et al., 2022), and AMPd-Up (Li et al., 2024), and used in vitro assays to characterize their antibacterial, hemolytic, and cytotoxic properties. In this study, we selected three lead AMPs (named TeBi1, TeRu4, and PeNi4) for further testing, due to their high antimicrobial activity and low toxicity. The three AMPs were tested in vivo as a prophylactic treatment prior to APEC inoculation (challenge trials). Immunomodulatory effects of these AMPs were tested in tissue harvested from animals involved in the challenge trials and in HD11 cells, a chicken macrophage-like cell line. Lastly, the AMPs were tested for their safety in a commercial production setting (pen trials). Overall, TeRu4 demonstrated activity in HD11 cell culture, while both TeBi1 and TeRu4 showed efficacy in challenge trials and maintaining safety in commercial production settings, supporting their potential for future translational studies.

## MATERIALS AND METHODS

### In Silico Peptide Selection

Candidate AMP sequences were predicted using three machine-learning-based software programs developed by our team: AMPlify, rAMPage, and AMPd-Up (Li et al., 2022, 2024; Lin et al., 2022). Candidate AMPs were evaluated for potential cytotoxicity using tAMPer (Ebrahimikondori et al., 2024), a machine-learning-based toxicity prediction tool, and those predicted to be cytotoxic were excluded from further evaluation.

### Peptide Synthesis

All AMPs for this study were purchased from GenScript (Piscataway, NJ, USA). AMP purity of ≥85% was verified using analytical HPLC. For in vitro assays and APEC challenge trials, AMPs were synthesized in 80 µg aliquots, while AMPs selected for pen trials were synthesized in 125, 250, or 500 mg aliquots.

### Bacterial Species and Growth

APEC strain EC317 (Allan et al., 1993) (clinical turkey isolate, obtained from VIDO-InterVac, Saskatchewan, CA), was used as the bacterial target for AMPs. Glycerol stocks were stored at -80°C. The primary streaks were done by plating the stocks on 5% Columbia blood agar (Thermo Fisher Scientific, Waltham, MA, USA) and incubating for 20-24 hours at 37°C under aerobic conditions, as described by Richter et al. (Richter et al., 2022). The secondary streaks were done by transferring 2-3 colonies from the primary streak on a new 5% Columbia blood agar plate. Colonies from the secondary streak were used for downstream analysis. Bacterial growth medium was prepared by dissolving 10.5 g Mueller Hinton Broth (Sigma-Aldrich, St. Louis. MO, USA) powder in 500 ml distilled water and autoclaving for sterility.

### In Vitro Lead Peptide Selection

Minimum inhibitory concentrations (**MICs**) of candidate AMPs were determined in vitro against EC317, using ASTs as previously described (Richter et al., 2022). For the selection process, candidate AMPs with MICs ≤ 32 µg/mL against EC317 were designated as hit AMPs and selected for further in vitro testing. Following the protocol by Richter et al. (Richter et al., 2022), hemolytic activities of hit AMPs were evaluated in porcine red blood cells (**RBCs**). Those with low hemolytic activity (HC_50_ ≥ 128 µg/mL, where HC_50_ is the AMP concentration that lyses 50% RBCs) were subsequently tested for cytotoxicity in human kidney HEK293 cells according to the protocol described (Richter et al., 2022). AMPs with CC_50_ ≥ 128 µg/mL (CC_50_, the AMP concentration that reduces cell viability to 50%) were designated as lead AMPs (Table 1).

**Table 1.**
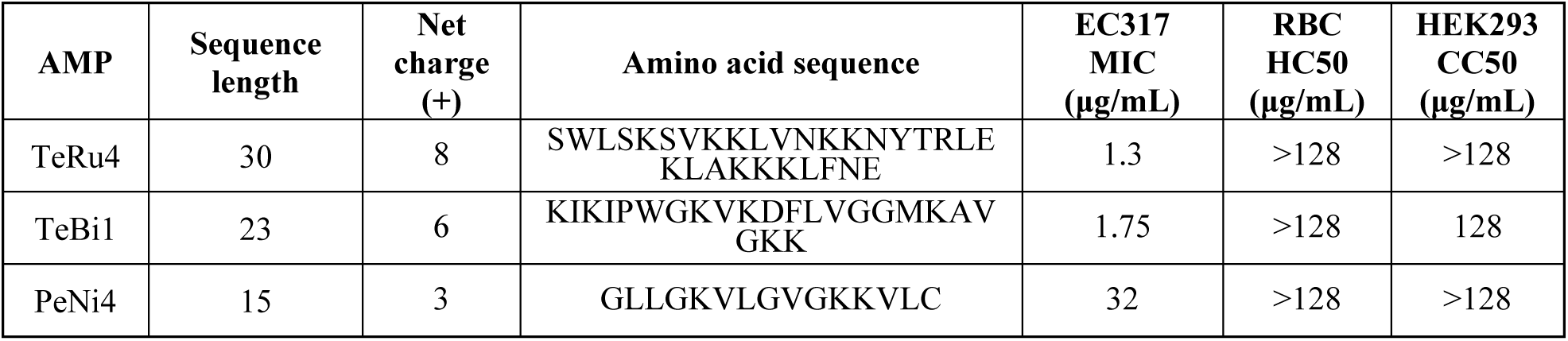
Physiochemical properties and in vitro activities of three lead antimicrobial peptides (AMP) TeRu4, TeBi1, and PeNi4. Listed for each AMP are their sequence length, net charge, and amino acid sequence. Each AMP was synthesized as the C-terminal carboxylic acid. In vitro activities include the antimicrobial activity against avian pathogenic *E. coli* (APEC) EC317 (minimum inhibitory concentration, MIC), the hemolytic activity against porcine red blood cells (RBCs) (hemolytic concentration 50%, HC50), and cytotoxicity towards HEK293 human cells (cytotoxic concentration 50%, CC50).

### Care and Use of Animals

All animal experiments were approved by the University of Saskatchewan and University of Victoria Animal Care Committees (Protocols #19940213 and AE-23-012, respectively) and conducted in accordance with the guidelines of the Canadian Council on Animal Care.

### APEC Challenge Trials

On day 18 of incubation (trial day -3), 50-57 fertilized Ross broiler chicken eggs per treatment were injected with either 1, 5, 10, or 20 μg of AMP, or PBS control. When the birds hatched on day 21 post-fertilization (trial day 0), they were neck-tagged with unique identification numbers and co-housed on Biofresh bedding (Early’s Farm and Garden, Saskatoon, Canada), with a light:dark cycle of 23L:1D. Egg hatchability rates and average bird weights were recorded for each treatment group. One day after hatch, the birds were challenged with 5 × 10^4^ cfu EC317.

Before inoculation, APEC was cultured overnight at 37 °C in LB broth (Difco, Becton Dickinson, Sparks, MD, USA) in a shaking incubator (Model No: 1570, Sheldon Manufacturing, Inc., Cornelius, OR, USA) at 240 rpm. This culture was sub-cultured to a pre-determined OD_600_ of 0.7, at which time the bacteria were harvested by centrifugation, resuspended in phosphate-buffered saline (**PBS**) at the desired concentration, and administered to the birds by yolk sac injection.

The birds were given food (Masterfeeds, Humboldt, SK, Canada) and water ad libitum. Four days after hatch, approximately half of the birds (between 8-17) were euthanized. The remaining birds were euthanized seven days after hatch. Birds were monitored two to four times per day for symptoms, and birds reaching a humane intervention point were euthanized. Only the birds euthanized on schedule on days four and seven were included in the analyses. Bird survival was also tracked.

All analyzed birds were weighed after death, and swabs were taken aseptically from the air sac and pericardium and streaked on MacConkey agar (Difco, Becton Dickinson, Sparks, MD, USA) for a semi-quantitative assessment of APEC growth. Plates were scored from 0-3, representing no, low, medium, and severe levels of growth, respectively. Samples from the air sacs, cecal tonsils, and spleens of birds euthanized on scheduled days were preserved in RNALater (Thermo Fisher Scientific) at –20 °C for analysis by quantitative PCR (**qPCR**).

### Immunomodulatory Activity Assays with HD11 Cells

The effects of AMPs on in vitro cytokine production were assessed using an HD11 chicken semi-adherent macrophage-like cell line (Obtained from VIDO-InterVac). HD11 cells were cultured in RPMI medium with 1X Glutamax and 10% Fetal Bovine Serum (Thermo Fisher Scientific) at 41°C (normal chicken body temperature) and 5% CO_2_ and were routinely passaged every two days.

#### Cell Plating

HD11 cells were plated for experimentation between passages three to twelve. Suspended cells were collected with the growth medium, and adherent cells were obtained after washing with PBS and trypsinization with 0.25% Trypsin-EDTA in PBS (both from Thermo Fisher Scientific). The cells were centrifuged at 200 g for seven minutes and the pellet was resuspended in fresh medium. Trypan Blue (Thermo Fisher Scientific) was added to a sample of cells and live cells were counted on a Hausser Scientific Bright-Line Phase Hemacytometer (Fisher Scientific, Hampton, NH, USA). HD11 cells were plated at 2.5 × 10^4^ cells per well on Nunclon Delta 96-well plates (Thermo Fisher Scientific) and incubated overnight.

#### HD11 Experiment Overview

In all experiments, HD11 growth medium was used to resuspend and dilute AMPs and other reagents, and all incubations were carried out at 41°C and 5% CO_2_. All experiments included a medium-only control and a lipopolysaccharide (**LPS**)-only control (final concentration of 25 ng/mL). In all experiments other than the AMP-only Screen, 8 μg/mL Polymyxin B sulfate (**PB**) was used as a positive control as it is known to attenuate the LPS-triggered inflammatory response (Schromm et al., 2021; Yang et al., 2024). Prior to treatment, the supernatant was removed from the wells, therefore only the adherent cell monolayer was used during the experiments.

Four experimental approaches were devised to investigate the immunomodulatory activity of AMPs. First, HD11 cells were treated solely with AMPs (AMP-Only Screen). Second, cells were treated with a range of AMP concentrations followed by treatment with LPS from *E. coli* O111:B4 (Sigma-Aldrich), to test if the AMPs could modulate the immune response triggered by LPS (AMP Preincubation Screen). If an AMP showed immunomodulatory activity in the AMP Preincubation Screen, it was then tested in the AMP Preincubation with Wash Step and the AMP and LPS Coaddition experiments.

#### AMP-Only Screen

HD11 cells were treated with 100 µL 16 µg/mL TeRu4, TeBi1, or PeNi4, and incubated for 6 h.

#### AMP Preincubation Screen

HD11 cells were treated with 100 μL of 1, 2, 4, 8, or 16 μg/mL TeRu4, TeBi1, or PeNi4, and incubated for 3 h. After the incubation, 100 μL LPS (50 ng/mL) was administered to the cells, resulting in a final concentration of 25 ng/mL LPS. The cells were incubated for an additional 3 h.

#### AMP Preincubation with Wash Step

The cells were treated identically to the AMP Preincubation Screen but with the addition of a wash step prior to the addition of LPS. Briefly, after the first 3 h incubation, the medium (containing the AMP or PB) was removed from the wells and the cell monolayer was washed three times using fresh medium. One hundred μL LPS (25 ng/mL) was administered, and the cells were incubated for an additional 3 h.

#### AMP and LPS Coaddition

AMP was resuspended and diluted to prepare a 2X concentration curve of 32, 16, 8, 4, and 2 μg/mL. PB and LPS were prepared at 2X concentrations of 16 μg/mL and 50 ng/mL, respectively. At regular time intervals, 2X AMP or 2X PB was added in equal volume to 2X LPS to yield a 1X mixture of AMP or PB, and LPS. Mixtures were vortexed for one min and incubated for 30 min prior to being administered to cells at regular time intervals in the order in which they were made. The cells were incubated for 3 h.

#### Cell Harvest and RNA Isolation

Supernatant was removed from HD11 cells after the final incubation. Buffer RLT from the miRNeasy Micro Kit (Qiagen, Mississauga, ON, Canada) with 2M DL-dithiothreitol (Sigma-Aldrich) was added to lyse adherent cells at a volume of 130 μL per well, with each technical replicate comprising two wells in the 96-well plate. HD11 cells in lysis buffer were disrupted by pipetting up and down, vortexed, and stored at -80 °C until RNA isolation. In total, three technical replicates per treatment were collected for RNA isolation and qPCR analysis.

### Commercial Pen Trials

Each pen trial examined the efficacy against ECM and the safety of a single AMP. Cobb 500 or Ross 308AP broiler chicken breeds were used in pen trials (Supplementary Table S1). All breeds were sourced from a commercial hatchery (Abbotsford, BC, Canada), and the specific breed used in each trial is summarized in Supplementary Table S1. For pen trials, birds were sourced from older breeder flocks (a different flock for each trial) and separated by sex and treatment into individual mini pens following hatching.

AMPs were solubilized in 400 mL sterile diluent (**SD**) bags, obtained from the hatchery, to reach a concentration of 0.4 or 0.2 mg/mL, except in the 20µg TeRu4 trial where PBS was used instead. An AMP-free SD bag was used for control. The AMP and control solutions were then taken to the hatchery, for in ovo injections on day 18 post-fertilization, where 50 µL AMP or SD solution was injected into the air sac of 1,680 to 2,352 eggs using the Embrex Inovoject injection machine (Zoetis Inc., Parsippany, NJ, USA). After the injections, eggs were incubated for three days. On the hatching day (day 0), pips, culls, late-stage embryo mortalities (deaths occurring between day 15 and 21 of incubation), post-hatch mortalities, and nonviable eggs/embryos (total of pips, culls, late-stage embryo mortalities, and post-hatch mortalities) were counted and recorded.

The birds were then transferred to SJ Ritchie Research Farms (Abbotsford, BC, Canada) where they were cloacally sexed by professionals and then separated by sex and treatment. A total of 24 mini pens, within six blocks, were set up. The birds were weighed individually, then 50 birds were placed into each mini pen according to their sex and treatment, with each block containing each of the four experimental groups in a random order. Birds were provided ad libitum access to water and an all-vegetable, raised without antibiotic (**RWA**) feed (Trouw Nutrition, Chilliwack, BC, Canada). A step-down lighting program was implemented: from day 0 to day 21, birds experienced 6 h darkness per 24 h cycle; from day 21 to 28, 4 h darkness; and from day 28 to the end of the trial, 1 h darkness per day. Number of birds in each mini-pen was reduced to 12 on day 10 to prevent overcrowding of the mini-pens. Birds were checked three times per day throughout the trial and were weighed individually on days 10, 21, and 35. Any mortality during the pen trial was noted.

### qPCR Analyses

Multiplex qPCR assays were designed and separately validated for three chicken tissues: air sac, cecal tonsil, and spleen; and for the HD11 macrophage-like cell line. Primers and probes were designed for the normalizer genes *ribosomal protein L8* (*rpl8*) and *ribosomal protein S10* (*rps10*); the cytokine genes *IL-1β, IL-6, IL-8, IL-10,* and *IFN-γ;* and the gene for the transcription factor *IRF-4*. The primer and probe sequences are reported in Supplementary Table S2.

RNA was extracted from tissues with the miRNeasy Tissue/Cells Advanced Mini Kit and Qiavac vacuum manifold (Qiagen). Tissue samples were homogenized in 450 μL buffer RLT reduced with 2M DTT, loaded on the RNeasy mini spin column, washed on the Qiavac vacuum manifold, and eluted in 50 μL RNAse-free water. RNA was extracted from HD11 cells using the using the QIAGEN miRNeasy Tissue/Cells Advanced Micro Kit, per the manufacturer’s instructions, except for the final elution, where 11.2 μL RNase-free water was added to the column, centrifuged, and the eluate was added back to the column and centrifuged again.

The concentration of RNA from tissue samples was measured by NanoDrop (NanoDrop 1000 3.8.1; Thermos Fisher Scientific) and diluted to a concentration of approximately 100 ng/μL before complementary DNA (**cDNA**) synthesis, while RNA from HD11 cells was used neat due to its low concentration. All RNA samples were converted to cDNA using the Applied Biosystems High-Capacity cDNA Reverse Transcription Kit (Thermo Fisher Scientific), per the manufacturer’s instructions.

cDNA from HD11 cells and chicken tissues was diluted 1/20 after synthesis and analysed by their respective qPCR assays with reaction conditions outlined in Supplementary Table S3. All samples and controls were run in technical quadruplicates, and all plates included a non-template (negative) control and inter-plate standard as a positive control. Standard deviations between the Cycle Threshold (**Ct**) values of technical replicates were computed. If any deviation was greater than 0.5 cycles, the outlier value in that set of replicates was removed. Normalizer stability was verified by BestKeeper (Pfaffl et al., 2004). Mean values between technical replicates were taken into the next stage of analysis. Data was normalised against the geometric mean of rpl8 and rps10, and fold change was calculated using the 2^-ΔΔCt^ method, as previously described (Livak and Schmittgen, 2001). Final relative fold changes for each sample and cytokine were determined by comparing to the relevant control: median of the PBS-treated birds from day 4 (for challenge trials), median of the PBS-treated female birds (for pen trials), or the mean of medium-treated HD11 cells (for cell culture experiments).

### Statistical Analyses

Statistical analyses of the qPCR results were conducted in R version 3.6.3 (R Core Team, 2024). The data were tested for normality by the Shapiro–Wilk and Levene’s tests. They were then subjected to pairwise analyses by the Kruskal-Wallis and Wilcoxon rank-sum tests for non-parametric data with a significance threshold of α = 0.1. For the challenge and pen trials, this significance cut off was selected due to the high variability of in vivo data. For the HD11 experiments, this threshold was appropriate as the smallest p-value achievable using a rank-sum test with n = 3 was approximately 0.08.

The rest of the challenge trial data was analyzed in R version 4.4.0 and 4.5.1(R Core Team, 2024). Hatchability rates of treated eggs compared to control eggs, as well as rates of bacterial detection in the birds’ air sacs and pericardia, were evaluated using the Fischer’s exact test. The presence or absence of bacteria detected in the birds’ air sacs and pericardia was also compared to AMP dose using a Cochrane-Armitage trend analysis on both days 4 and 7. Cumulative weight gain of birds euthanized on days four and seven was evaluated using a Wilcoxon rank-sum test, and survival was graphed and evaluated using the log-rank test and the “survminer” package (Kassambara et al., 2016). Other challenge trial graphs were constructed using the packages “tidyverse,” “readr,” “janitor,” “superb,” “ggsignif,” “rstatix,”, “DescTools”, and “ggpubr” (Signorell, 2014; Wickham et al., 2015b; a, 2019; Kassambara, 2016, 2019; Firke, 2016; Ahlmann-Eltze and Patil, 2017; Cousineau, 2021).

An additional analysis was conducted to compare the relative cytokine abundance in bird air sacs to the air sac bacterial load scores of the same birds. The difference in cytokine abundance for each bacterial level (light, moderate, severe) was compared to the cytokine abundance for no growth using a Wilcoxon rank-sum test, and additionally, a correlation coefficient was determined using Spearman’s Rho.

Statistical analyses of hatchability, survival, mean weights, feed conversion ratio, and flock uniformity in pen trials were conducted using R (version 4.1.3) and the packages: “survival”, “survminer”, “openxlsx”, “readxl”, “tidyverse, “zoo”, “car”, “ggplot2”, “ggsignif”, and “MHTdiscrete” (Fox et al., 2001; Therneau, 2001; Zeileis et al., 2004; Wickham et al., 2007, 2019; Schauberger and Walker, 2014; Wickham and Bryan, 2015; Kassambara et al., 2016; Yalin Zhu, Wenge Guo, 2016; Ahlmann-Eltze and Patil, 2017). For pen trials, statistical comparisons between treatment and control groups used Wilcoxon rank-sum tests (hatchability and production parameters) and log-rank tests (survival), with production and survival analyzed separately by sex. P-values were corrected for multiple comparisons using the Šidák method.

## RESULTS

### In Vitro Peptide Selection

The 875 candidate AMPs identified by rAMPage (Lin et al., 2022), AMPlify (Li et al., 2022), and AMPd-Up (Li et al., 2024) were previously tested for antimicrobial activity and cellular toxicity in vitro (Richter et al., 2022). Of these, 331 with MICs ≤ 32 µg/mL against EC317 were designated as hit AMPs. Among hit AMPs, 275 exhibited minimal hemolytic activity (HC_50_ ≥ 128 µg/mL) against porcine RBCs. Of these, 62 also demonstrated minimal cytotoxicity (CC_50_ ≥ 128 µg/mL) in HEK293 cells. These were classified as lead AMPs. Three representative lead AMPs (TeRu4, TeBi1, and PeNi4) were selected for further experiments. TeRu4 and TeBi1 demonstrated lower MIC values than PeNi4. The physiochemical properties of these AMPs and their in vitro activities are summarized in Table 1. These AMPs were evaluated in vitro using HD11 cells and in vivo using APEC challenge trials and pen trials.

### qPCR Primer Design and Validation

The sequences of the qPCR primers and probes, reaction conditions, and assay efficiencies are reported in Supplementary Tables S2-4. Primer efficiencies ranged from 93-108% in the air sac, 92-107% in the cecal tonsil, 87-107% in the spleen, and 84-101% in HD11 cells.

### APEC Challenge Trials

To assess the prophylactic efficacy of TeBi1, TeRu4, and PeNi4 in chickens, fertilized eggs were injected in ovo at day 18 of incubation with either 1, 5, 10, or 20 μg AMP, or a PBS control. One day after hatch, the birds were challenged with 5 × 10^4^ cfu APEC strain EC317 by yolk sac injection.

Egg hatchability was between 90-98% in all treatment groups, with no significant differences between treated birds and control birds (data not shown). Notably, when birds were treated with 20 μg AMP (Supplementary Figure S1), treated birds appeared to die faster than untreated birds within the first few days post-infection, though there were no significant differences in overall survival. The spleens of birds treated with 20 μg AMP were analyzed by qPCR, and some differences were observed, suggesting that the peptides were bioactive, but the qPCR results were not conclusive (data not shown), and subsequent challenge trials tested the AMPs on a dose curve at 1, 5, or 10 μg. In the latter trials, there were no significant differences in survival probability with any dose of AMP (≤10 μg) compared to control (Supplementary Figure S2).

Birds treated with TeBi1 had significantly higher weights on day 7 than control birds (Figure 1a). Birds treated with 1 or 5 μg of TeBi1 were approximately 25% heavier than control birds (p = 0.03 and p = 0.07, respectively), and birds treated with 10 μg TeBi1 were approximately 50% heavier than control birds (p = 2 × 10^-4^). There were no significant differences in weight gain in birds treated with TeRu4 or PeNi4 (Supplementary Figure S3).

**Figure 1:**
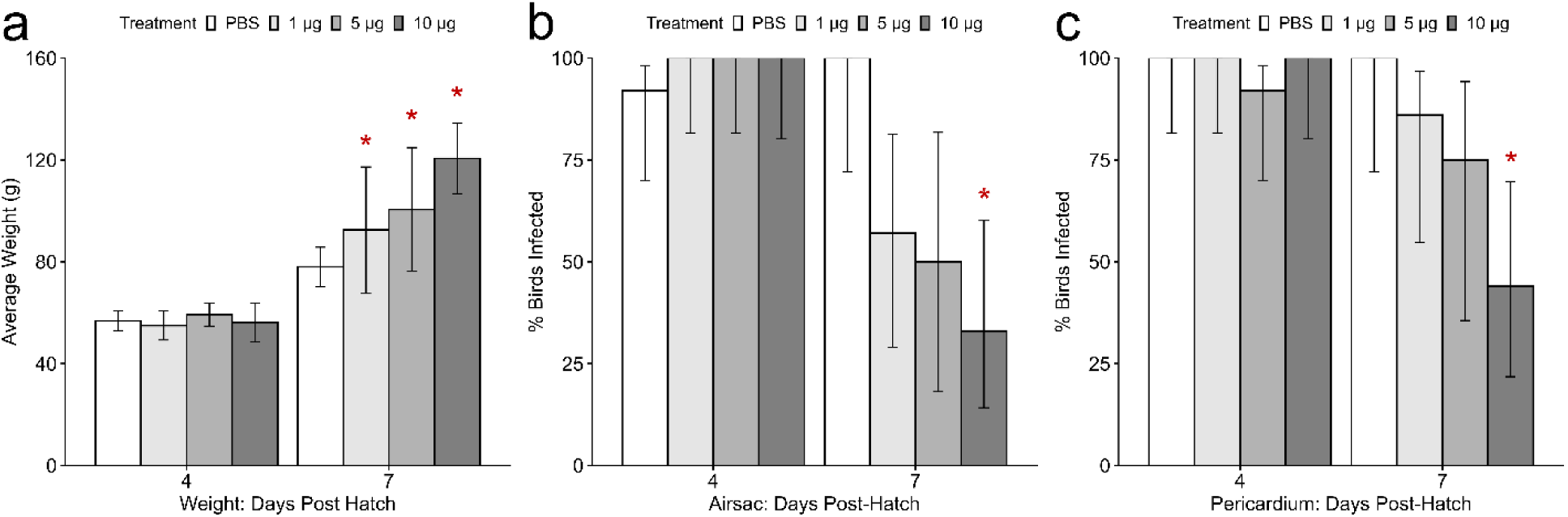
a) Weight gain of TeBi1 treated birds between day 4 and 7. Birds, euthanized on schedule, were weighed individually, and values shown are median ± median absolute deviation. On day 7, birds treated with 1 or 5 μg TeBi1 were approximately 25% heavier than control birds, and birds treated with 10 μg TeBi1 were approximately 50% heavier than control birds. Significance (p ≤ 0.05) relative to the phosphate buffered saline (PBS) control is indicated by an asterisk. b/c) Percent of TeBi1 treated birds infected by avian pathogenic *E. coli* (APEC) at on day 4 and 7 post-hatch in the b) air sacs and c) pericardia. Birds were euthanized as scheduled and aseptic swabs were taken from the tissues and streaked onto MacConkey agar. The percentage of birds in each treatment group that resulted in bacterial growth is shown, with the error bars denoting a 90% Wilson confidence interval. Each treatment group was compared to the PBS control by Fisher’s Exact Test. Significance (p ≤ 0.05) relative to the PBS control is indicated by an asterisk. On day 7, there were significantly fewer birds with detectable bacteria in the 10 μg TeBi1 treatment group than the control group, in both tissues.

On day 7, birds treated with 10 μg TeBi1 had significantly lower rates of infection in both the air sac (p = 0.01) (Figure 1b) and pericardium (p = 0.03) (Figure 1c). Increased peptide dose was also generally correlated with decreased numbers of birds with detectable bacteria in either the air sac or the pericardium (p = 0.02 and p = 0.008, respectively, also on day 7). There were no significant differences in bacterial detection rates in the air sacs and pericardia of birds treated with TeRu4 or PeNi4 (Supplementary Figure S4), but for TeRu4-treated birds on day 4, there was a slight but similar trend of decreased numbers of infected birds with increasing AMP dose (air sac: p = 0.009, pericardium: p = 0.1), and for PeNi4, on day 4, in the air sac only, there was a slight trend of increasing numbers of infected birds with increased AMP dose (p = 0.1).

Increased bacterial load in the air sac was correlated with increased levels of transcript abundance of certain cytokines in the air sac (Figure 2). Across all three challenge trials, birds with light, moderate or heavy levels of active APEC infection in their air sacs had significantly higher relative abundance of *IL-1β* transcripts (p = 2 × 10^-4^, 1 × 10^-6^, and 7 × 10^-13^, respectively) than birds with no bacterial growth in their air sacs (Figure 2a). They are also positively correlated by Spearman’s rho, with ρ = 0.58 and p = 1 × 10^-19^. The same trend, though weaker, is observed with *IL-8*. While ρ = 0.32 (p = 3 × 10^-6^) overall, birds with light, moderate, or heavy levels of APEC infection also had significantly higher relative abundance of *IL-8* transcripts (p = 0.03, 3 × 10^-6^, and 5 × 10^-6^, respectively) than birds with no bacterial growth in their air sacs (Figure 2b).

**Figure 2.**
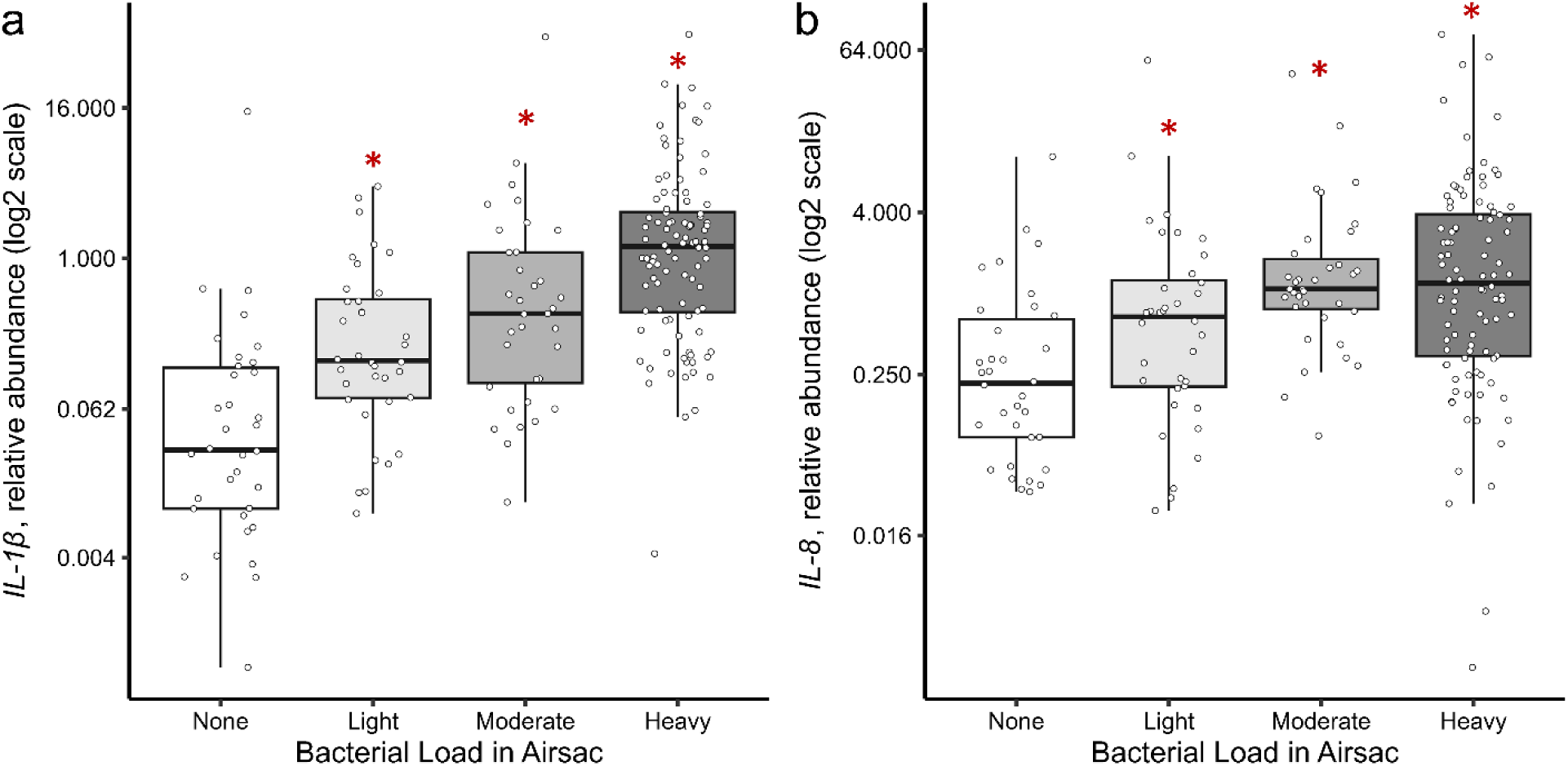
Relative transcript abundance of a) *Interleukin (IL)-1β* and b) *IL-8* in the air sacs of birds treated with ≤10 μg TeRu4, TeBi1, or PeNi4. Each data point represents the air sac of a single bird. The light, moderate, and heavy infection groups were compared to the non-infected group (none) using the Wilcoxon rank-sum test. Significance (p ≤ 0.05) relative to non-infected group is indicated by an asterisk.

In the cecal tonsil (Supplementary Figure S5), qPCR results show minimal changes due to treatment with PeNi4, with only a 2-fold increase in the inflammatory cytokine *IL-8* in the 10 μg group on day 4 (p = 0.1); and with TeRu4, a 2-fold decrease in *IL-8* in the 1 μg group on day 4 (p = 0.1). In contrast, there are numerous changes due to treatment with TeBi1, including a 2.4-fold increase in *IL-8* transcripts in the 5 μg group on day 4 (p = 0.1), and a 5-fold decrease in *IL-8* transcripts in the 10 μg group on day 7 (p = 0.06). *IFN-γ* expression increased 1.4-fold (p = 0.03) and 2.5-fold (p = 0.03) in the 5 and 10 μg groups, respectively, on day 4, but decreased 1.4-fold (p = 0.03) and 1.1-fold (p = 0.09) in the 1 and 10 μg groups, respectively, on day 7. With respect to *IL-1β,* on day 7, the TeBi1 5 μg group showed a 5-fold decrease (p = 0.07). For the anti-inflammatory transcript *IL-10,* there is a 2.5-fold decrease (p = 0.07) with 5 μg TeBi1 on day 7. Finally, TeBi1 treatment was also associated with changes in *IRF-4* transcripts: the 10 μg group was associated with a 1.7-fold increase on day 4 (p = 0.04), and the 5 μg group was associated with a 1.5-fold increase on day 7 (p = 0.1).

In the spleen (Supplementary Figure S6), fewer responses are observed. For TeRu4, there is a 2-fold decrease in *IL-8* transcripts on day 7 (p = 0.1) after treatment with 1 μg. For PeNi4, there is a 5-fold increase in *IL-10* transcripts on day 7 (p = 0.09) after treatment with 5 μg. For TeBi1, while there are no differences in the inflammatory cytokines *IFN-γ, IL1β,* or *IL-8*, there is a 3-fold increase in *IL-10* transcripts on day 4 (p = 0.03) after treatment with 10 μg. There are also multiple increases in *IRF-4* transcripts: on day 4, a 2-fold increase after treatment with 1 μg (p = 0.1), a 3-fold increase after treatment with 5 μg (p = 0.1); and on day 7, a 1.3-fold decrease after treatment with 1 μg (p = 0.1).

In the air sac (Figure 3), all observed changes are on day 7, after treatment with 10 μg TeBi1 relative to the PBS control. There is a 10-fold decrease in *IFN-γ* transcripts (p = 0.1), a 3-fold decrease in *IL-1β* transcripts (p = 0.07), and a 5-fold decrease in *IL-8* transcripts (p = 0.02). Additionally, there is a 50-fold decrease in *IL-10* transcripts (p = 0.1), and a 10-fold decrease in *IRF-4* transcripts (p = 0.1).

**Figure 3.**
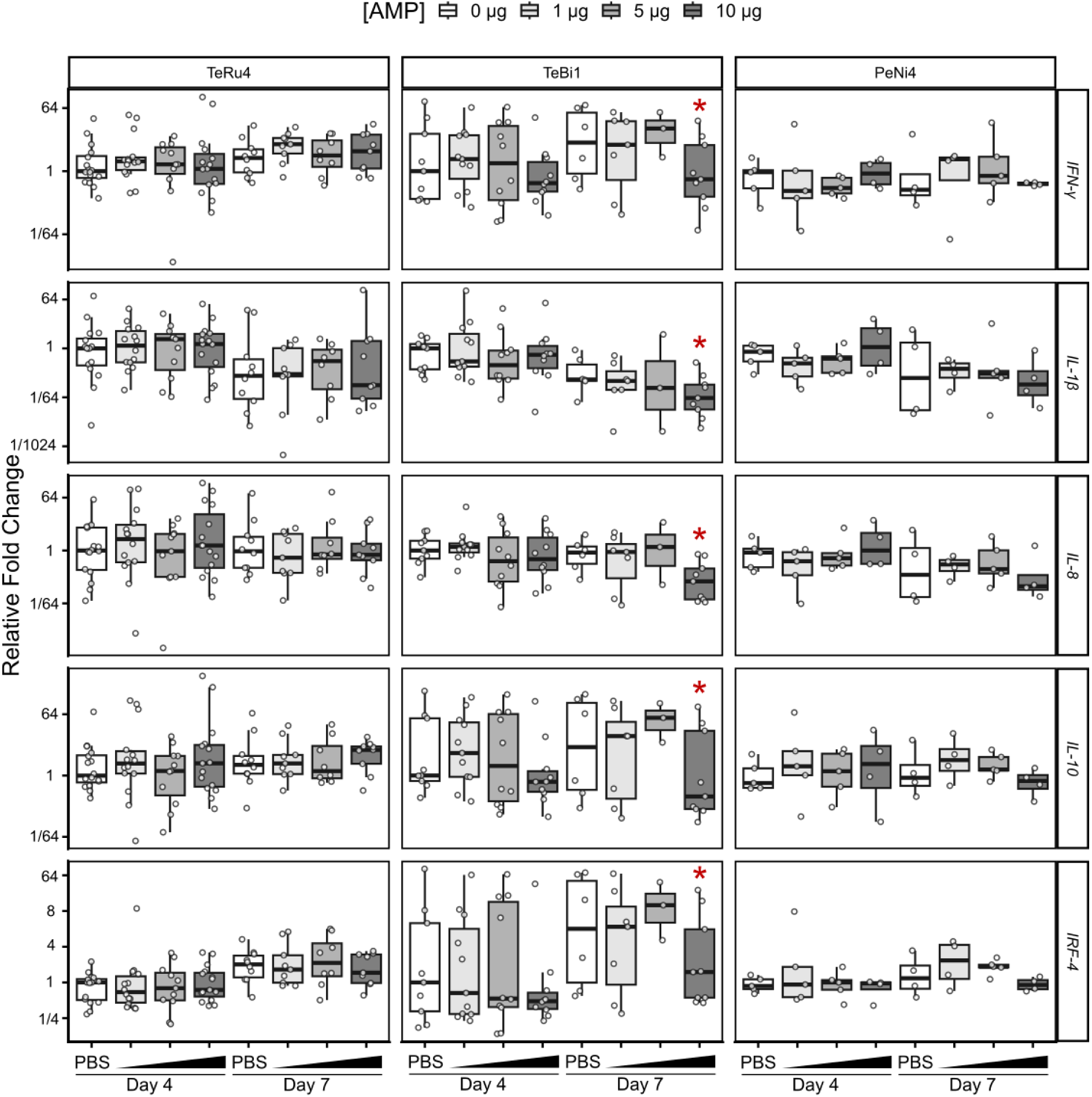
Relative fold change values of cytokine transcripts in the air sac of avian pathogenic *E. coli* (APEC) challenged birds treated before hatch with 1, 5, or 10 μg antimicrobial peptide (AMP). Birds were euthanized on day 4 or 7 after hatch, and all significance comparisons are relative to the phosphate-buffered saline (PBS) on their respective days. Asterisks indicate significance at p ≤ 0.1. *IFN*, interferon; *IL,* interleukin; *IRF*, interferon regulatory factor.

### HD11 Experiments

In addition to challenge trials, the ability of TeRu4, TeBi1, and PeNi4 to influence the cellular immune response was tested in an HD11 cell model system. In all HD11 experiments (n=21), the LPS induction control resulted in a wide-ranging, but consistent increase in relative fold change compared to the medium control for *IL-1β* (14 - 677), *IL-8* (8 - 249), and *IL-6* (55 - 3,284) (Figure 4 and Supplementary Figures S7 and S8). With the exception of the AMP-only experiment, all stated fold-changes are relative to the LPS induction control. The induction of cytokine transcripts by LPS was significantly attenuated by PB in all but a single cytokine measurement from one experiment (n=15), with fold change decreases compared to LPS alone ranging from 6-166 for *IL-1β*, 2-34 for *IL-8*, and 11-335 for *IL-6* (Figure 4 and Supplementary Figure S8). The preincubation, preincubation with wash step, and coaddition experiments were performed in biological triplicate, and the AMP-only experiments in biological duplicate. While only one representative biological replicate is shown per experiment, these selected replicates capture the overall trends in fold changes observed across replicates (unless otherwise stated). The fold change values stated are that of the representative replicate.

**Figure 4.**
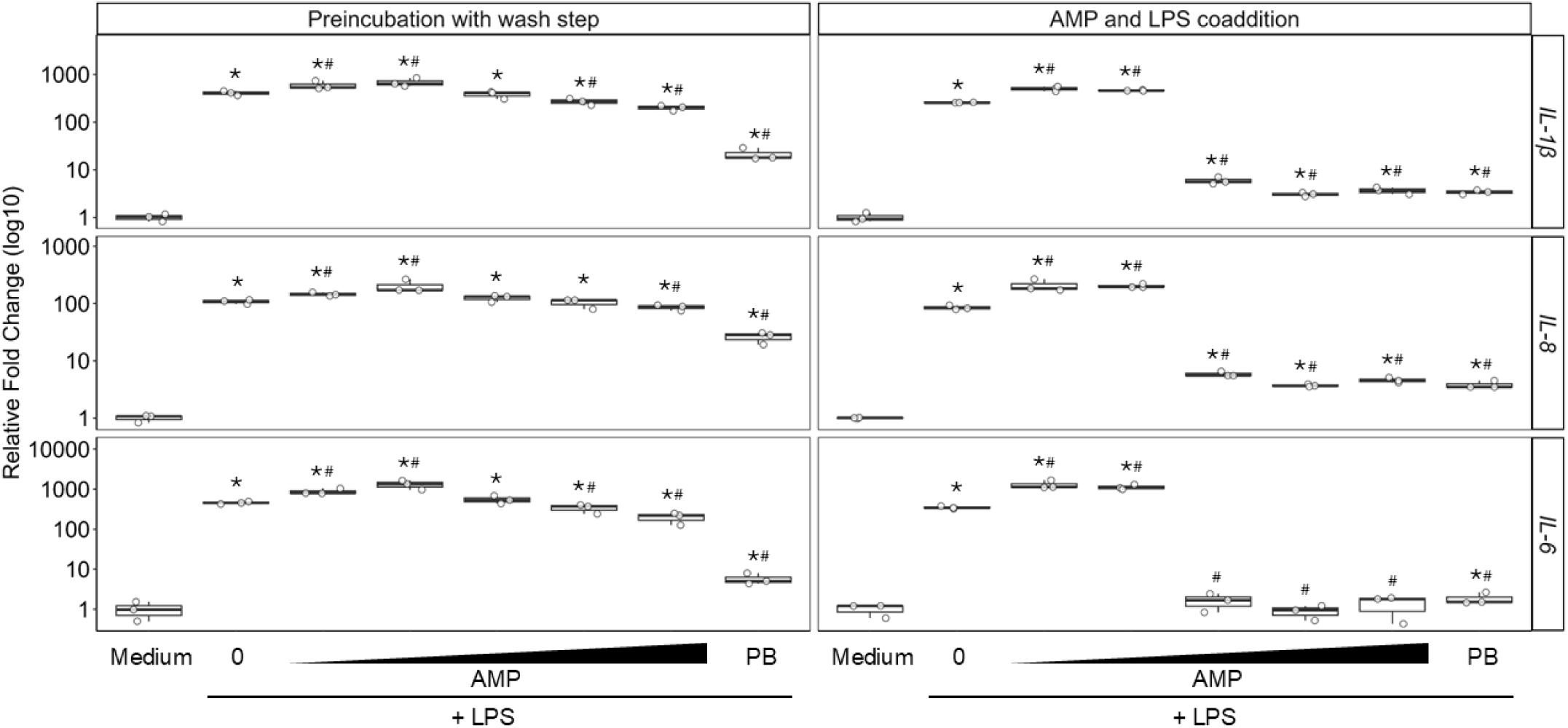
Relative fold change of *Interleukin (IL)-1β*, *IL-8*, and *IL-6* from HD11 cells as determined by qPCR in the “preincubation with wash step” (left column) and “AMP and LPS coaddition” experiments (right column). In the preincubation with wash step experiment, cells were incubated for 3 h with medium, a range of 2-fold serially diluted antimicrobial peptide (AMP; 1 to 16 µg/mL TeRu4), or 8 µg/mL polymyxin B sulfate (PB), followed by three washes prior to the addition of 25 ng/mL lipopolysaccharide (LPS) for an additional 3 h. In the coaddition experiment, cells were incubated for 3 h with medium, or with 25 ng/mL LPS mixed with a range of 2-fold serially diluted AMP (1 to 16 µg/mL TeRu4), or 8 µg/mL PB. A significant difference from the medium control is indicated by “*”, and “#” indicates a significant difference from LPS treatment (p-value < 0.1, Mann-Whitney-U test). Open circles represent the fold change of one technical replicate relative to the medium control. The horizontal lines represent the median of the three technical replicates within each treatment condition, the boxes show the interquartile range, and the vertical whiskers represent the minimum and maximum values. Shown are representative experiments of three biological replicates.

#### AMP-Only Screen of TeRu4, TeBi1, and PeNi4

To assess the ability of AMPs alone to induce cytokine production in HD11 cells, cells were incubated with 16 µg/mL of each AMP or 8 µg/mL PB and cytokine levels were compared to a medium-only control. Treatment with TeRu4 resulted in a 2-fold increase in *IL-1β* (in one of two biological replicates) and 3-fold increase in *IL-8,* while TeBi1 lead to a 4-fold decrease in *IL-6* (Supplementary Figure S7). Treatment with PeNi4 did not significantly alter the abundance of any of the three cytokines, while TeBi1 had no significant effect on *IL-1β* and *IL-8,* and TeRu4 had no significant effect on *IL-6* (Supplementary Figure S7). Treatment with PB alone did not significantly or consistently change *IL-1β, IL-8*, and *IL-6* levels (Supplementary Figure S7).

#### Preincubation Screen of TeRu4, TeBi1, and PeNi4

To examine if AMP affect HD11 response to LPS, cells were preincubated with AMP or PB (positive control), followed by treatment with LPS. Here and in subsequent experiments, medium only and LPS only served as negative and positive controls, respectively. A 2-fold decrease was observed for *IL-1β* and *IL-6* following preincubation with 8 µg/mL TeRu4 (Supplementary Figure S8). Preincubation with 16 µg/mL TeRu4 led to a significant reduction in *IL-1β, IL-8*, and *IL-6* levels by 13-fold, 4-fold, and 28-fold, respectively, (Supplementary Figure S8). Similarly, treatment with 8 µg/mL PB significantly decreased *IL-1β*, *IL-8* (in 2 of 3 biological replicates), and *IL-6* abundance by 10-fold, 3-fold, and 20-fold, respectively (Supplementary Figure S8). While a significant 4-fold increase was observed for *IL-8* at 4 µg/mL TeRu4, this observation was only seen in two of three biological replicates (Supplementary Figure S8). Preincubation with TeBi1 and PeNi4 did not consistently induce a significant change in *IL-1β, IL-6*, and *IL-8* transcript levels across biological replicates (Supplementary Figure S8). The attenuation of LPS-induced responses of *IL-1β, IL-8*, and *IL-6* transcripts after TeRu4 preincubation prompted further investigation via the subsequent experiments to provide some insight into possible mechanisms of action.

#### Preincubation Experiment with Wash Step

The effects on HD11 cells of transient exposure to TeRu4, followed by its removal and the addition of LPS, was examined. A significant decrease in *IL-1β, IL-8,* and *IL-6* transcripts remained after preincubation with 8 µg/mL PB (21-, 5-, and 80-fold respectively) and 16 µg/mL TeRu4 (2-, 1-, and 2-fold, respectively) (Figure 4). Preincubation with 8 µg/mL TeRu4 induced a significant decrease in *IL-1β* levels by 2-fold and *IL-6* by 1-fold in two of three biological replicates (Figure 4). Unlike in the previous preincubation experiment (without a wash step), treatment with 1 and 2 µg/mL TeRu4 followed by the wash step resulted in a 1- and 2-fold increase respectively in *IL-1β* and *IL-8* transcript levels(but not consistently in all biological replicates) (Figure 4). For *IL-6* in this experiment, treatment with 1 and 2 µg/mL TeRu4 resulted in a2-and 3-fold increase respectively across all three biological replicates (Figure 4).

#### Coincubation of TeRu4 and LPS

The impact of co-incubating TeRu4 and LPS prior to treating HD11 cells on cytokine transcript expression was assessed. Coaddition of LPS with either PB or TeRu4 (at concentrations ranging from 4 to 16 µg/mL) resulted in significant decreases in all cytokine transcripts across all three biological replicates (Figure 10). Specifically, co-treatment with PB and LPS led to reductions of 75-fold for *IL-1β*, 24-fold for *IL-8*, and 226-fold for *IL-6* (Figure 4). Similarly, co-treatment with 16 µg/mL TeRu4 and LPS decreased *IL-1β* abundance by 70-fold, 18-fold for *IL-8*, and 191-fold for *IL-6* (Figure 4). LPS and 8 µg/mL TeRu4 reduced *IL-1β* transcripts by 83-fold, *IL-8* by 23-fold, and *IL-6* by 363-fold (Figure 4). LPS and 4 µg/mL TeRu4 resulted in reductions of 46-fold for *IL-1β*, 15-fold for *IL-8*, and 205-fold for *IL-6* (Figure 4). For *IL-6*, 4-16 µg/mL TeRu4 repressed the LPS induction so much that expression levels were not significantly different from the medium only negative control (observed in two of three replicates at 4 µg/mL) (Figure 4). Like what was observed in the previous experiment, 1 and 2 µg/mL of TeRu4 resulted in significantly increased transcript abundance in *IL-1β* (2-fold), *IL-8* (2-fold), and *IL-6* (3-fold) relative to the LPS control, but unlike the previous experiment, these results were consistent across three biological replicates (Figure 4).

### Commercial Pen Trials

Pen trials were designed to test the efficacy against ECM and the safety of treating chickens with TeBi1, TeRu4, and PeNi4 in a commercial production setting. Late-stage viability (viable embryos from day 15 of incubation until placement in pens) was between 88 and 98% in all pen trials. No statistically significant differences were detected between the treatment and control groups (Supplementary Figure S9). There were no significant differences between the treatment and control groups for post-hatch mortality, culls, pips, and late embryo mortality.

For the first 10 days, there were no statistical differences between the survival probability of males and females (Supplementary Figure S10). Thus, the sexes were combined for subsequent analyses. No significant differences in survival probability were detected between treatment and control groups for any AMPs during the first 10 days of the pen trials, indicating that the tested AMPs had no measurable impact on ECM (Supplementary Figure S11). However, for the full duration of the pen trials (35 days), sex-specific differences in survival probabilities were observed for TeRu4 at 20 µg and TeBi1 at 10 µg (Supplementary Figure S12). For these two pen trials, the sexes were kept separate for analysis (Figure 5, Supplementary Figure S13). Notably, TeRu4 at 20 µg significantly increased the survival probability of female birds compared to the controls by approximately 4.9 % by day 35 (p = 0.04). Moreover, TeBi1 at 20 µg significantly increased the survival probability of the birds compared to the control by approximately 4.4% by day 35 (p = 0.02). There was no other difference in survival probabilities between the treatment and control groups by day 35. In the 10 µg TeRu4 and TeBi1 trials, survival curves for both sexes closely overlapped (Supplementary Figure S13).

**Figure 5.**
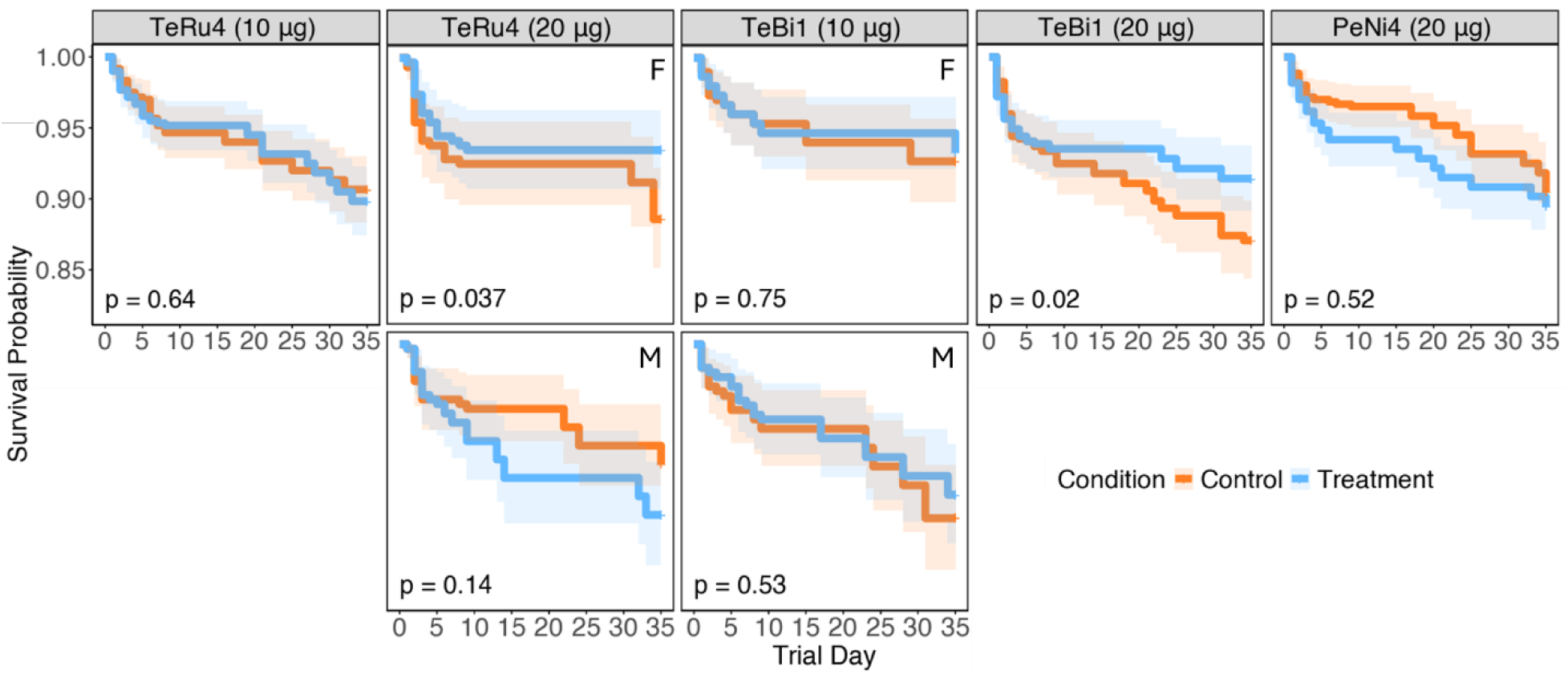
Kaplan-Meier curves depicting the survival probabilities of experimental groups during pen trials across different antimicrobial peptides (AMP) and dosages. Columns represent AMPs at specific dosages and each AMP–dosage combination corresponds to a separate pen trial. Sex-specific differences in survival probabilities were observed in the TeRu4 (20 µg) and TeBi1 (10 µg) trials. For these trials, males and females were analyzed separately, and each panel is labeled with “F” for females or “M” for males in the top right corner to indicate the sex of the birds included in the analysis. P-values, as determined by Log-rank tests, are displayed in the bottom-left corner of each panel.

Significant differences in mean weight between treatment and control groups were observed only in the 20 µg TeBi1 trial (Supplementary Figure S14). Notably, on day 0, mean weights in the treatment groups were approximately 2.8% lower in males and 2.4% lower in females compared to controls (p = 0.02 and 0.03). However, on day 10, no significant difference in mean weights were observed. Moreover, in the 10 µg TeRu4 trial, numerous birds of both sexes exhibited lower body weights on days 21 and 35, particularly in the treatment group, although this difference was not statistically significant. This observation may suggest subclinical effects of TeRu4 at lower doses, high biological variability among individual birds, or environmental effects, all of which warrant further investigation. By day 35, no significant differences in mean weight were detected (Supplementary Figure S14).

No statistically significant differences were detected between FCR of the treatment and control groups on day 35, suggesting that the tested AMPs did not exhibit measurable growth-promoting properties (Supplementary Figure S15). Of interest, there were occasions where FCR values were below 1 (Supplementary Figure S15). One was observed between days 0 and 10, in the 10 µg TeBi1 trial (FCR = 0.997). Further, in the 20 µg TeRu4 trial, two FCR values were below 1 (FCR = 0.923 and 0.961) in the second interval, between days 11 and 21. Overall, the mean FCR values were consistently the lowest in the first interval, and the highest in the last interval, between days 21 and 35, for both sexes. These findings suggest that AMPs did not significantly alter FCR under the tested conditions, and observations during the first interval of certain pen trials warrant further investigation.

Coefficient of variation (**CV**) percentages were calculated as a measure of flock uniformity. There was no significant difference in the CV percentages of the treatment and control groups on day 35 (Supplementary Figure S16). Among males, on day 10 in the 20 µg TeBi1 trial, the treatment group had an approximately 2.8% lower CV compared to the control group, but this difference did not reach statistical significance (p = 0.07) (Supplementary Figure S16). Moreover, the birds in the 10 µg TeRu4 trial exhibited greater variability in flock uniformity on days 21 and 35, with CV values exhibiting greater range than those in the control, consistent with the presence of birds with low body weight.

The qPCR results were analyzed with sexes separated, and significant differences are reported for both comparisons between birds in the same sex group that received different treatments, and for birds in the same treatment group with differences between sexes. As there were some significant differences in transcript abundance between male and female birds in the same treatment groups, the results are shown with sexes separated in Figure 6.

**Figure 6.**
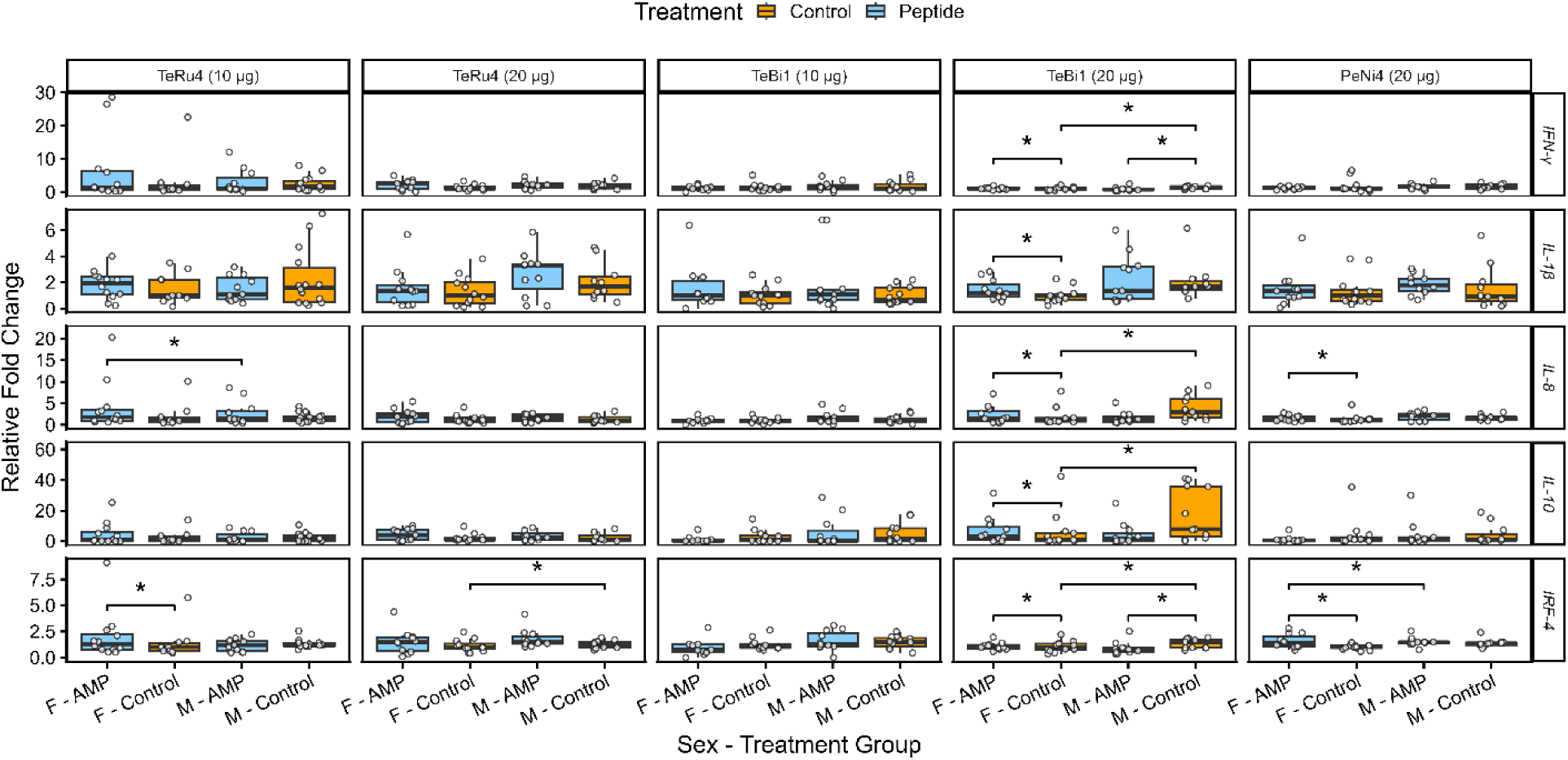
Relative fold change values of cytokine transcripts in the spleens of pen trial birds (no pathogen challenge) treated with 10 or 20 μg TeRu4, TeBi1, or PeNi4. Birds were euthanized 10 days after hatch. Asterisks indicate significance at p ≤ 0.1. F, female birds; M, male birds. Additional details are available in the Figure 5 legend.

After treatment with 10 μg of TeRu4, female birds had a 1.8-fold increase (p = 0.1) in *IFN-γ* transcripts relative to (female) control birds, and in that trial, male controls birds had a 1.2-fold increase (p = 0.1) in *IRF-4* transcripts relative to female control birds. After treatment with 20 μg of TeRu4, male birds had a 1.3-fold increase (p = 0.08) in *IRF-4* transcripts relative to control birds.

After treatment with 20 μg PeNi4, female birds had a 1.4-fold increase (p = 0.02) in *IRF-4* transcripts relative to control birds. Additionally, in that trial, male control birds had a 1.3-fold increase (p = 0.01) in *IRF-4* transcripts and a 1.6-fold increase (p = 0.04) in *IL-8* transcripts relative to female control birds.

When considering the sexes separately, there were no significant differences in transcript abundance in birds treated with 10 μg TeBi1 (Figure 6). As there were no significant differences between sexes in this trial, qPCR results from that trial were also combined (Supplementary Figure S17). Here, a 1.2-fold increase (p = 0.1) in *IL-1β* transcripts was observed. Examination of the transcript abundances in birds treated with 20 μg TeBi1 revealed numerous differences relative to PBS controls.

After treatment with 20 μg TeBi1, male birds had a 2-fold decrease in *IFN-γ* and *IRF-4* transcripts (p = 0.02 for both), and a 3-fold decrease in *IL-10* and *IL-8* transcripts (p = 0.03 and 0.02, respectively) relative to control birds. In that trial, male control birds had higher abundance of all tested transcripts than female control birds: 2-fold more *IFN-γ* transcripts (p = 0.1), 3-fold more *IL-10* transcripts (p = 0.06), 1.7-fold more *IL-1β* transcripts (p = 0.008), 3-fold more *IL-8* transcripts (p = 0.01), and 2-fold more *IRF-4* transcripts (p = 0.1). Additionally, male birds treated with 20 μg TeBi1 had significantly lower abundance of certain transcripts than female birds in the same treatment group, a 1.3-fold decrease in both *IFN-γ* and *IRF-4* transcripts (p = 0.09 for both).

## DISCUSSION

This study demonstrates that antimicrobial peptides can improve early health outcomes in broiler chickens while maintaining production performance under commercial conditions. Across complementary in vivo and in vitro assays, two of the three lead AMPs, TeBi1 and TeRu4, showed efficacy and immunomodulatory activity consistent with their intended roles as antibiotic alternatives for preventing APEC-associated diseases. Here, AI-assisted discovery is relevant for enabling large-scale candidate generation and triage that would be impractical by hand. All outcome data from the in vivo and in vitro assays were analyzed using nonparametric methods, Kruskal–Wallis for omnibus comparisons and Wilcoxon rank-sum for pairwise contrasts. Given the inherent variability of in vivo data and the exploratory nature of this study, a significance threshold of α = 0.1 was used, as specified in the methods.

First, in controlled challenge trials, pre-hatch administration of TeBi1 was associated with reduced post-hatch bacterial burden and improved early growth. At 10 μg per egg, TeBi1 significantly lowered culture detection of APEC in the air sac and pericardium by day 7 and yielded an approximate 50% increase in body weight relative to controls at the same time point. These findings support a direct antibacterial contribution in vivo and align with TeBi1’s favorable in vitro profile (Lin et al., 2022). Concordant decreases in pro-inflammatory cytokine transcripts in air sac tissue at day 7 (Figure 3) further suggest that limiting bacterial load mitigates downstream inflammatory responses. In contrast, TeRu4 and PeNi4 did not reproducibly reduce organ-specific bacterial detection in the challenge setting at tested doses up to 10 μg, indicating peptide-specific differences in in vivo antibacterial activity despite their performance in in vitro experiments. Consistent with this, positive correlations between air sac bacterial loads and *IL-1β* (ρ = 0.58) and *IL-8* (ρ = 0.32) expression (Figure 2) both support the validity of the infection model and indicate that AMP-mediated reductions in bacterial burden translate to dampened inflammatory signaling.

Second, the HD11 macrophage-like cell experiments demonstrate the potential for TeRu4 to attenuate the inflammatory effects of LPS. Preincubation with TeRu4 and its coaddition with LPS elicited robust and dose-dependent attenuation of *IL-1β*, *IL-6*, and *IL-8* transcripts, with effects at higher exposures of TeRu4 that were comparable in direction and magnitude to PB. While the persistence of attenuation throughout the experiments (Figure 4) may indicate that TeRu4 can interact both with LPS and with HD11 cells to mitigate inflammation, further study is required to confirm these mechanisms. Notably, a biphasic pattern was observed: attenuation of *IL-1β, IL-6,* and *IL-8* at higher exposures of TeRu4, but mild potentiation at sub-threshold doses. Taken together, these data support an immunomodulatory profile for TeRu4 that could help contain excessive inflammation induced by LPS, even when direct antibacterial effects are limited in vivo.

Third, pen trials provide critical evidence of field-relevant safety and performance. In ovo administration of TeBi1 or TeRu4 at 10 or 20 μg per egg did not adversely affect hatchability, FCR, or flock uniformity through day 35. Importantly, survival probabilities improved modestly but significantly in two settings: a 4.9% increase in females with 20 μg TeRu4 and a 4.4% increase in all birds with 20 μg TeBi1 by day 35. Although effect sizes were small, they are operationally meaningful at commercial scale and were achieved without detectable adverse effects in growth or efficiency. These findings support a favorable benefit-risk profile for both peptides when delivered through standard in ovo automation.

Together, the in vivo and in vitro data outline complementary roles for these leads. TeBi1 appears to act primarily through direct antibacterial activity in the immediate post-hatch period, reducing tissue colonization and the associated inflammatory response. TeRu4 shows consistent anti-inflammatory activity in a macrophage model and over 35 days translates to a survival advantage in females at higher dose, suggesting that immunomodulation can contribute to resilience under farm conditions. These sex-specific responses may reflect differences in immune system maturation rates or hormone-immune interactions, as male broilers typically show different growth and immune development patterns than females (Leitner et al., 1989; Younis et al., 2023). The patterns observed in both survival and cytokine transcript baselines warrant additional investigation into sex-dependent pharmacodynamics (**PD**), immune set-points, and potential interactions with breeder age and genetics.

While the results presented here are encouraging, this study also has several limitations. First, the challenge trials used a single APEC strain and a prophylactic in ovo regimen. Broader pathogen panels, additional exposure routes, and therapeutic or peri-hatch dosing schedules should be evaluated to define the generalizability and optimal use cases. Second, although this study shows safety and performance neutrality in pens, pharmacokinetics (**PK**), peptide stability in the embryonic and early post-hatch milieu, and residue depletion profiles remain to be established. Third, gene expression endpoints were measured at discrete time points and in a single cell line or selected tissues. High-resolution temporal profiling, including protein-level cytokine measurements and cellular phenotyping in multiple cell lines and tissues, would sharpen mechanistic inference and help explain biphasic responses at lower TeRu4 concentrations. Further, some FCR values below 1.0 in early trial periods likely reflect measurement artifacts related to initial hydration status rather than true feed conversion, highlighting the importance of standardized weighing protocols in early post-hatch periods. Finally, while the analyses used conservative non-parametric tests and multiple-comparison corrections, some findings were near the chosen α threshold of 0.1, reflecting expected variability in vivo and underscoring the need for replication across flocks and sites.

The results have practical implications for poultry health programs. The in ovo delivery method offers multiple advantages: it integrates cleanly into existing hatchery workflows, avoids post-hatch handling, and ensures uniform exposure across flocks. This delivery approach, combined with a demonstrated safety profile, minimizes barriers to commercial adoption. From an economic perspective, even modest survival improvements translate to substantial numbers at commercial scale, while early bacterial control reduces the need for subsequent therapeutic interventions. The complementary mechanisms of TeBi1 (direct antibacterial) and TeRu4 (immunomodulatory) suggest potential for combination strategies that leverage both activities. Future work should benchmark these AMPs against current animal husbandry standards and evaluate alternative delivery formats, such as water supplementation during high-risk periods, to optimize protection throughout the production cycle.

In conclusion, AI-discovered AMPs, TeBi1 and TeRu4 show complementary efficacy and safety signals in broilers when administered in ovo. TeBi1 reduces early APEC burden and inflammation and improves early growth, whereas TeRu4 attenuates HD11 inflammatory responses and improves survival under commercial conditions without compromising performance. These findings support advancement to dose-optimization, PK and residue studies, expanded pathogen panels, and multi-site validation, with an eye toward integrated antimicrobial stewardship in poultry production.

## DECLARATION OF AI AND AI-ASSISTED TECHNOLOGIES IN THE WRITING PROCESS

During the preparation of this work, the authors used ChatGPT and Gemini to improve grammar, readability, and language. After using this tool/service, the author(s) reviewed and edited the content as needed and takes full responsibility for the content of the publication.

## ACKNOWLEDGMENTS

The authors thank Genome Canada and Genome British Columbia for financial support through the large scale applied projects program (#291PEP) to Birol I., Helbing C.C., Hoang L.M.N, Fraser E., and Hof F. Additional funding was provided by Chicken Farmers of Canada, Canadian Hatching Egg Producers, Investment Agriculture Foundation of British Columbia (INV106, INV247), and Amphoraxe Life Sciences Inc. Partial support for trainees were provided by Mitacs (IT28945, IT43363), University of Victoria, University of British Columbia, and Amphoraxe Life Sciences Inc.

