## Supplemental Tables and Figures for "Efficacy and safety evaluation of artificial intelligence-identified antimicrobial peptides for use against avian pathogenic *Escherichia coli* in the poultry industry"

[†] BC Cancer, Vancouver, BC V5Z 1L3, Canada
[‡] British Columbia Centre for Disease Control, Public Health Laboratory, Vancouver, BC V6Z R4R, Canada
[§] Genome Science and Technology Graduate Program, University of British Columbia, Vancouver, BC V6T 1Z4, Canada
[#] Department of Biochemistry and Microbiology, University of Victoria, Victoria, BC, V8P 5C2, Canada
[‖] Canada's Michael Smith Genome Sciences Centre at BC Cancer, Vancouver, BC V5Z 4S6, Canada
[††] Amphoraxe Life Sciences Inc., BC V6L 3C9, Canada
[‡‡] SJ Ritchie Research Farms, Abbotsford, BC V4X 2N4 Canada
[§§] Vaccine and Infectious Disease Organization (VIDO), University of Saskatchewan, SK, S7N 5E3, Canada
[##] Communicable Disease Prevention and Response, British Columbia Centre for Disease Control, Vancouver, BC V6Z R4R, Canada
[※] School of Population and Public Health, The University of British Columbia, Vancouver, BC, V6T 1Z3, Canada
[⁑] Department of Pathology and Laboratory Medicine, University of British Columbia, Vancouver, BC V6T 1Z4, Canada
[⁂] Department of Chemistry, University of Victoria, Victoria, BC, V8P 5C2, Canada
[⸎] Department of Medical Genetics, University of British Columbia, Vancouver, BC V6T 1Z3, Canada

[*]These authors contributed equally to this work.

**TABLE OF CONTENTS**


| | |
|---|---|
| **Supplementary Table S1** | AMP and dosage tested on two breeds of chicken |
| **Supplementary Table S2** | qPCR assay primer and probe sequences |
| **Supplementary Table S3** | qPCR reaction conditions |
| **Supplementary Table S4** | qPCR assay efficiencies |
| **Supplementary Figure S1** | Survival plots of chickens treated with 20 µg AMP |
| **Supplementary Figure S2** | Survival plots of chickens treated with 1, 5, or 10 µg AMP |
| **Supplementary Figure S3** | Weight gain of chickens treated with TeRu4 and PeNi4 between days 4 and 7 |
| **Supplementary Figure S4** | Viable bacterial detection rates in the air sacs and pericardia of birds treated with TeRu4 and PeNi4 |
| **Supplementary Figure S5** | Relative fold change of cytokine transcripts in the cecal tonsil of birds treated with 1, 5, or 10 µg AMP |
| **Supplementary Figure S6** | Relative fold change of cytokine transcripts in the spleen of birds treated with 1, 5, or 10 µg AMP |
| **Supplementary Figure S7** | Relative fold change values of cytokine transcripts in HD11 cells treated with AMP |
| **Supplementary Figure S8** | Relative fold change values of cytokine transcripts in LPS-treated HD11 cells after preincubation with AMP |
| **Supplementary Figure S9** | Raw percentage values for hatchability parameters across various AMPs and dosages |
| **Supplementary Figure S10** | Survival plots of male and female birds during the first 10 days of the pen trials |
| **Supplementary Figure S11** | Survival plots of treatment and control birds during the first 10 days of the pen trials |
| **Supplementary Figure S12** | Survival plots of male and female birds on days of the pen trials |
| **Supplementary Figure S13** | Survival plots of birds on day 35 of the pen trials |
| **Supplementary Figure S14** | Distribution of mean weights of birds during pen trials |
| **Supplementary Figure S15** | Feed conversion ratio (FCR) of birds during pen trials |
| **Supplementary Figure S16** | Flock uniformity of birds during pen trials across |
| **Supplementary Figure S17** | Relative fold change values of cytokine transcripts in the spleens of pen trial birds |

### SUPPLEMENTARY TABLES

**Supplementary Table S1.** Antimicrobial peptide (AMP) and dosage tested on two chicken breeds.

| AMP and Dosage Tested | Chicken Breed Used |
|---|---|
| TeBi1 - 10 µg | Ross 308AP |
| TeBi1 - 20 µg (Hatch Only) | Ross 308AP |
| TeBi1 - 20 µg | COBB 500 |
| TeRu4 - 10 µg | COBB 500 |
| TeRu4 - 20 µg | Ross 308AP |
| TeRu4 - 20 µg (Hatch Only) | COBB 500 |
| PeNi4 - 20 µg | COBB 500 |

**Supplementary Table S2.** qPCR assay primer/probe sequences. All sequences are 5' → 3'. Cy5, Cy5 fluorophore; FAM, FAM fluorophore; HEX, fluorophore; IABkFQ, Iowa Black FQ modified quencher; IAbRQSp, Iowa Black Rq Sp quencher; *IFN*, interferon; *IL,* interleukin; *IRF*, interferon regulatory factor; Mplex, multiplex reaction; *rps*, ribosomal protein small subunit encoding; TAO, TAO internal quencher; ZEN, ZEN internal quencher.

| Assay Group | Gene Target | Forward Primer | Reverse Primer | Probe |
|---|---|---|---|---|
| Mplex1/4 | *IL-1β* | CTTCGACATCAACCAGAA | CGACATGTAGAGCTTGTA | FAM-TGCTTCGTG-ZEN-CTGGAGTCACC-IABkFQ |
| | *rpl-8* | CAACCATCAGGAGAGATG | CTGGGTCTTTAGTTTTCC | HEX-TCGGTCTCA-ZEN-TTGCTGCTCGG-IABkFQ |
| | *rps-10* | GCTCGTTGATAAGAATGT | CAAATTGCTCTTTCACATA | Cy5-AGCCATGCA-TAO-GTCTCTGAAATCC-IAbRQSp |
| Mplex2/5 | *IL-8* | GTAGGACGCTGGTAAAGA | GGTGGATGAACTTAGAATGA | FAM-ACTGGCACC-ZEN-GCAGCTCATTC-IABkFQ |
| | *IL-10* | CCAGGTCAAAGAGAGTTA | CAGCAGCCTTTAAATCAA | HEX-TCCTATTAG-ZEN-AACCAACTGCTTAACTGCT-IABkFQ |
| Individual | *IFN-γ* | GTGGACCTATTATTGTAGAGA | GATGCTGAAGAGTTCATTC | FAM-CGCTGGATT-ZEN-CTCAAGTCGTTCAT-IABkFQ |
| Individual | *IL-6* | AGTGCCTGAATGTTTTAG | ACCGTAAGAAATGTAACAG | Cy5-CAATCCTCT-TAO-GTTACCAATCTGCCAC-IAbRQSp |
| Individual | *IRF-4* | AGGTGACAACTTCTAGTC | CTCAGCAGCTTTTCTATG | FAM-CTTCCTCTG-ZEN-GCTGTTATCCTCTGG-IABkFQ |

**Supplementary Table S3.** qPCR reaction conditions. All assays use hot-start enzymes and are subject to an initial denaturation consisting of 9 minutes at 95 °C. All assays run for 45 cycles. Qiacuity Probe (QIAcuity Probe PCR Kit; Qiagen, Mississauga, ON, Canada); Sensifast Probe (Sensifast Probe No-ROX kit; FroggaBio, Concord, ON, Canada); see Supplementary Table S2 for other definitions.

| Tissue/Cell | Assays | Thermocycle temperature sequence (°C) | Time sequence (s) | Enzyme Mix | Primer[1]/Probe Concentration (nM) |
|---|---|---|---|---|---|
| Air sac, cecal tonsil, spleen | Mplex1, Mplex2, *IFN-γ, IRF-4* | 95, 56.5, 72 | 45, 30, 45 | Qiacuity Probe | 300/100 |
| HD11 cell line | Mplex4, Mplex5, *IL-6* | 95, 60, 72 | 15, 30, 30 | Sensifast Probe | 700/100 |

[1]Per primer

**Supplementary Table S4.** qPCR assay efficiencies by assay and tissue type. See supplementary table S2 for definitions.

| Tissue | Assay Group | Gene Target | Efficiency |
|---|---|---|---|
| Air sac | Mplex1 | *IL-1β* | 92.82% |
| | | *rpl-8* | 102.52% |
| | | *rps-10* | 102.70% |
| | Mplex2 | *IL-8* | 107.52% |
| | | *IL-10* | 107.86% |
| | Individual | *IFN-γ* | 97.51% |
| | Individual | *IRF-4* | 98.01% |
| Cecal tonsil | Mplex1 | *IL-1β* | 93.69% |
| | | *rpl-8* | 98.01% |
| | | *rps-10* | 95.41% |
| | Mplex2 | *IL-8* | 102.70% |
| | | *IL-10* | 101.78% |
| | Individual | *IFN-γ* | 91.99% |
| | Individual | *IRF-4* | 106.86% |
| Spleen | Mplex1 | *IL-1β* | 91.50% |
| | | *rpl-8* | 106.86% |
| | | *rps-10* | 99.42% |
| | Mplex2 | *IL-8* | 90.98% |
| | | *IL-10* | 103.00% |
| | Individual | *IFN-γ* | 86.88% |
| | Individual | *IRF-4* | 90.44% |
| HD11 cells | Mplex4 | *IL-1β* | 84.32% |
| | | *rpl-8* | 86.08% |
| | | *rps-10* | 88.94% |
| | Mplex5 | *IL-8* | 100.80% |
| | | *IL-10* | 98.44% |
| | Individual | *IL-6* | 91.05% |

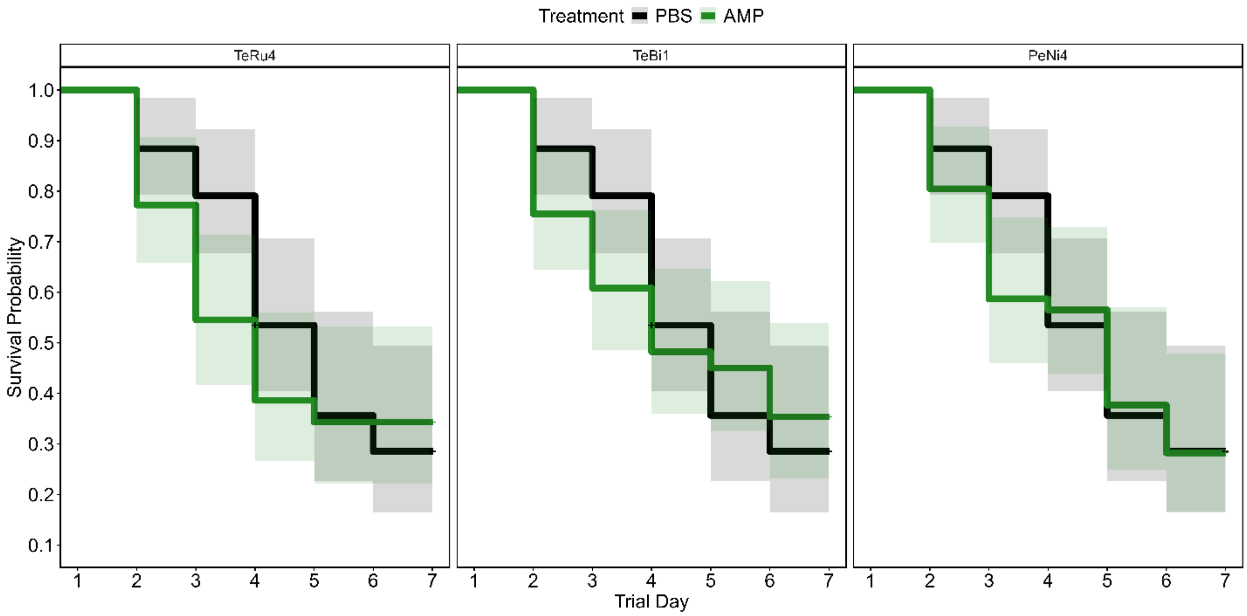


**Supplementary Figure S1.** Kaplan-Meier survival plots of chickens treated with 20 µg TeRu4, TeBi1, or PeNi4. There were no significant differences in survival between treatment groups.

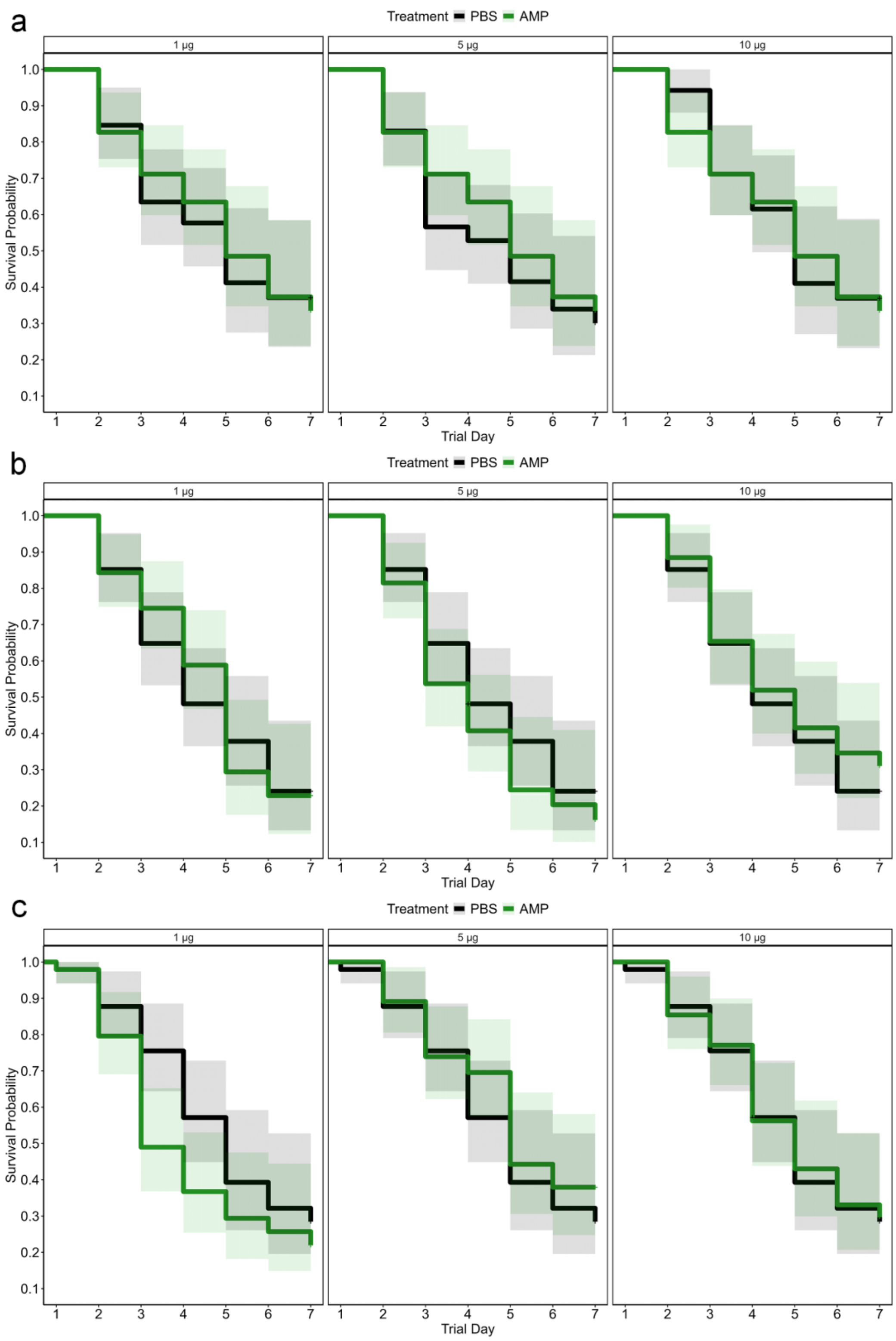
a
Treatment PBS AMP
1 µg
5 µg
10 µg
Survival Probability
Trial Day
b
Treatment PBS AMP
1 µg
5 µg
10 µg
Survival Probability
Trial Day
c
Treatment PBS AMP
1 µg
5 µg
10 µg
Survival Probability
Trial Day

**Supplementary Figure S2.** Kaplan-Meier survival plots of birds treated with either 1, 5, or 10 µg antimicrobial peptide (AMP). Plots shown are for a) TeRu4, b) TeBi1, or c) PeNi4. There were no significant differences in survival between treatment groups.

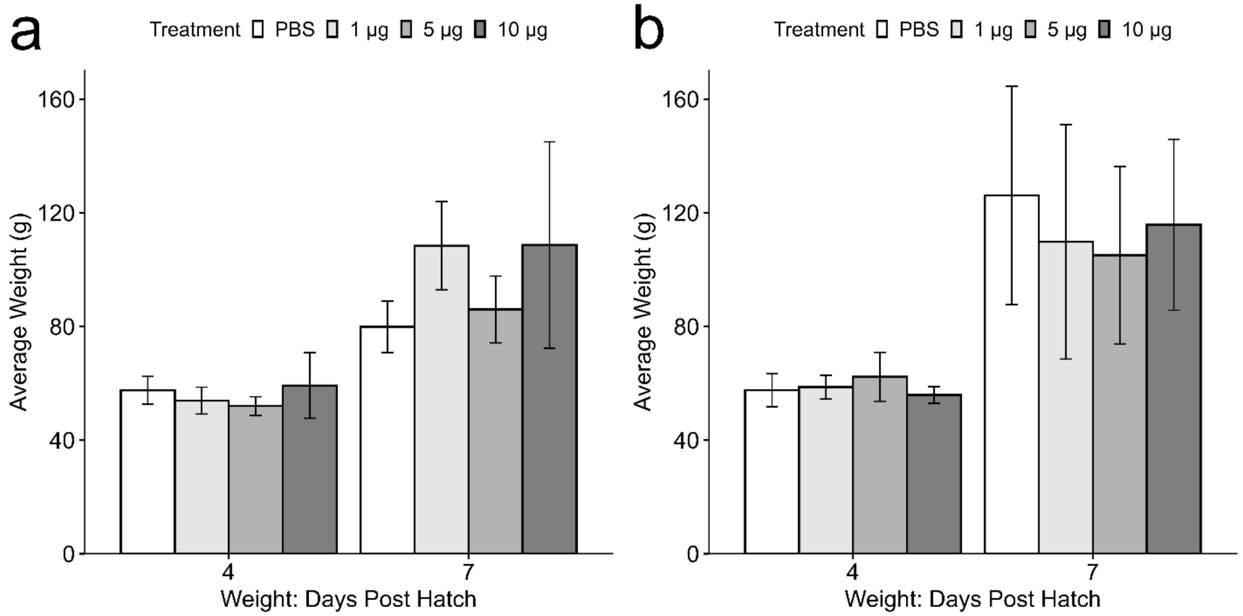


**Supplementary Figure S3.** Weight of birds treated with different doses of either a) TeRu4 and b) PeNi4 from day 4 to day 7 compared to their respective phosphate buffered saline (PBS) controls. Birds were weighed individually and the values shown are median ± median absolute deviation (MAD). There were no significant differences in bird weight with treatment.

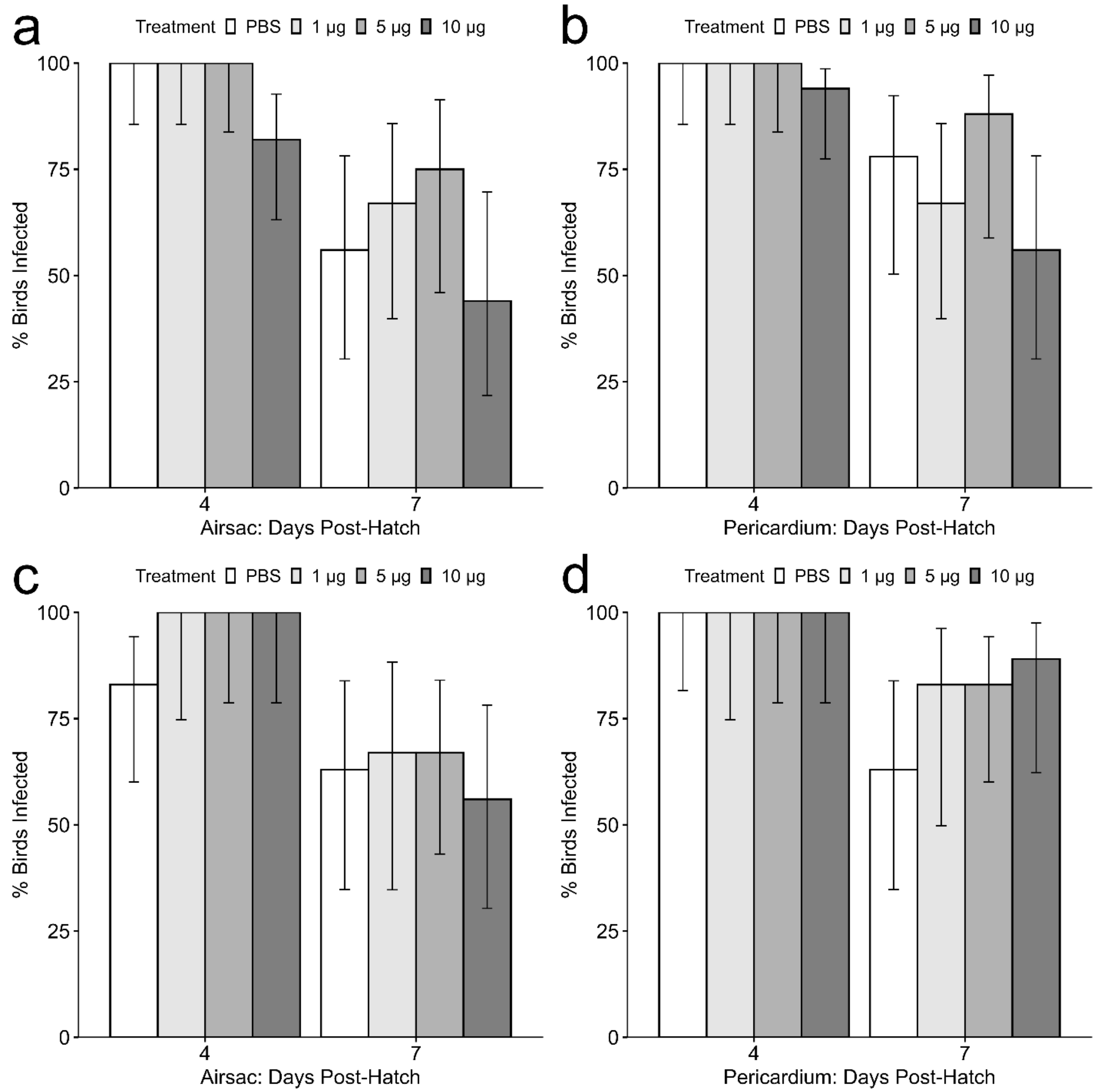


**Supplementary Figure S4.** Percent of birds infected by avian pathogenic *E. coli* (APEC) at specific time points post-hatch in the air sacs and pericardia of birds treated with 1, 5, or 10 µg TeRu4 and PeNi4 or phosphate buffered saline (PBS) control: a) TeRu4, air sac; b) TeRu4, pericardium; c) PeNi4, air sac; d) PeNi4, pericardium. Birds were euthanized as scheduled and aseptic tissue swabs were streaked onto MacConkey agar. The percentage of swabs in each treatment group that resulted in bacterial growth is shown, with the error bars denoting a 90%

Wilson confidence interval. Each treatment group was compared to the PBS control by Fisher's Exact Test. No significant differences were observed between treatments.

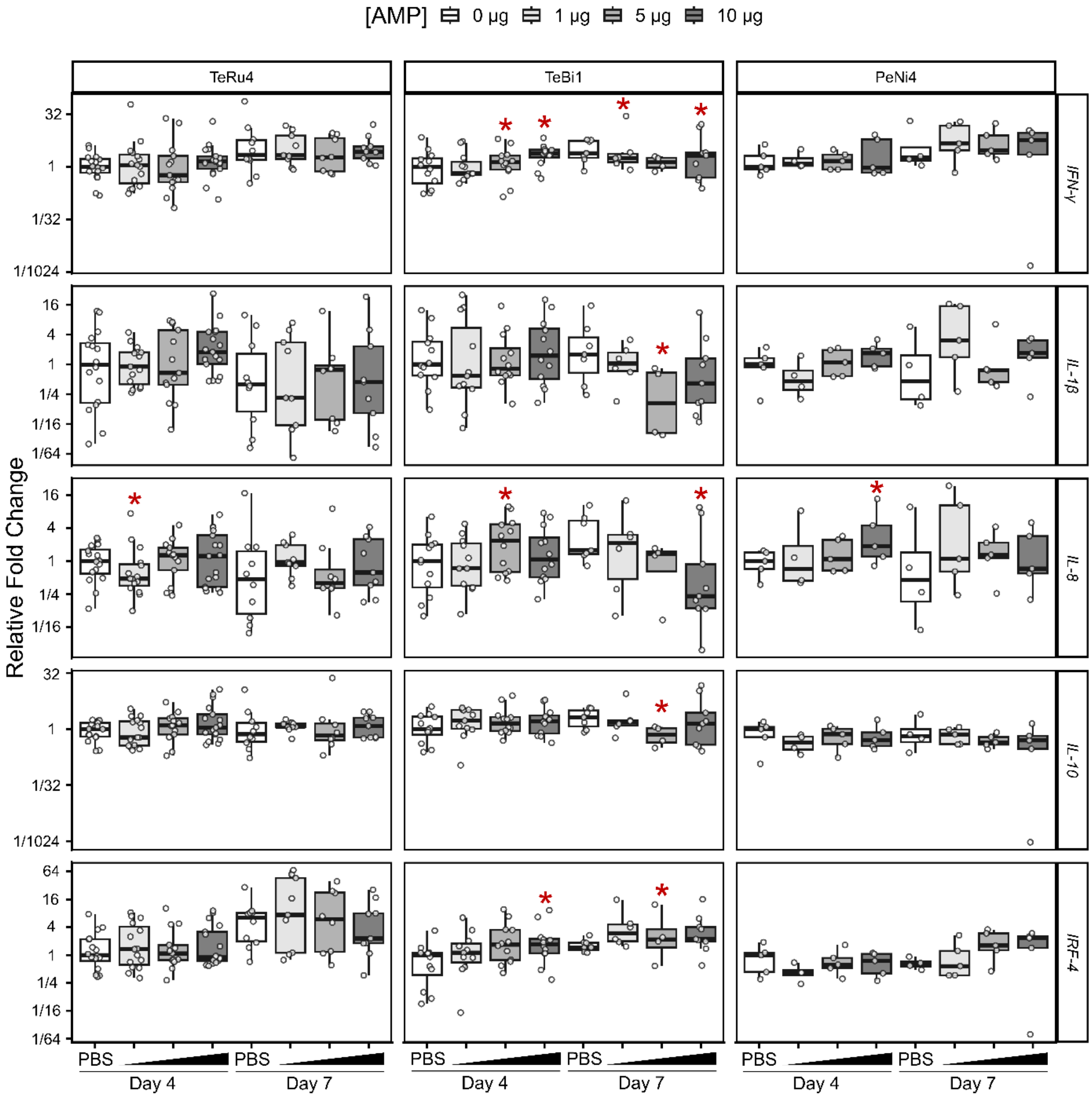


**Supplementary Figure S5.** Relative fold change values of cytokine transcripts in the cecal tonsil of avian pathogenic *E. coli* (APEC) challenged birds treated before hatch with 1, 5, or 10 μg antimicrobial peptide (AMP). Birds were euthanized on day 4 or 7 after hatch, and all values are relative to the PBS median on their respective days. Asterisks indicate significance at $p \leq 0.1$. *IFN*, interferon; *IL,* interleukin; *IRF*, interferon regulatory factor.

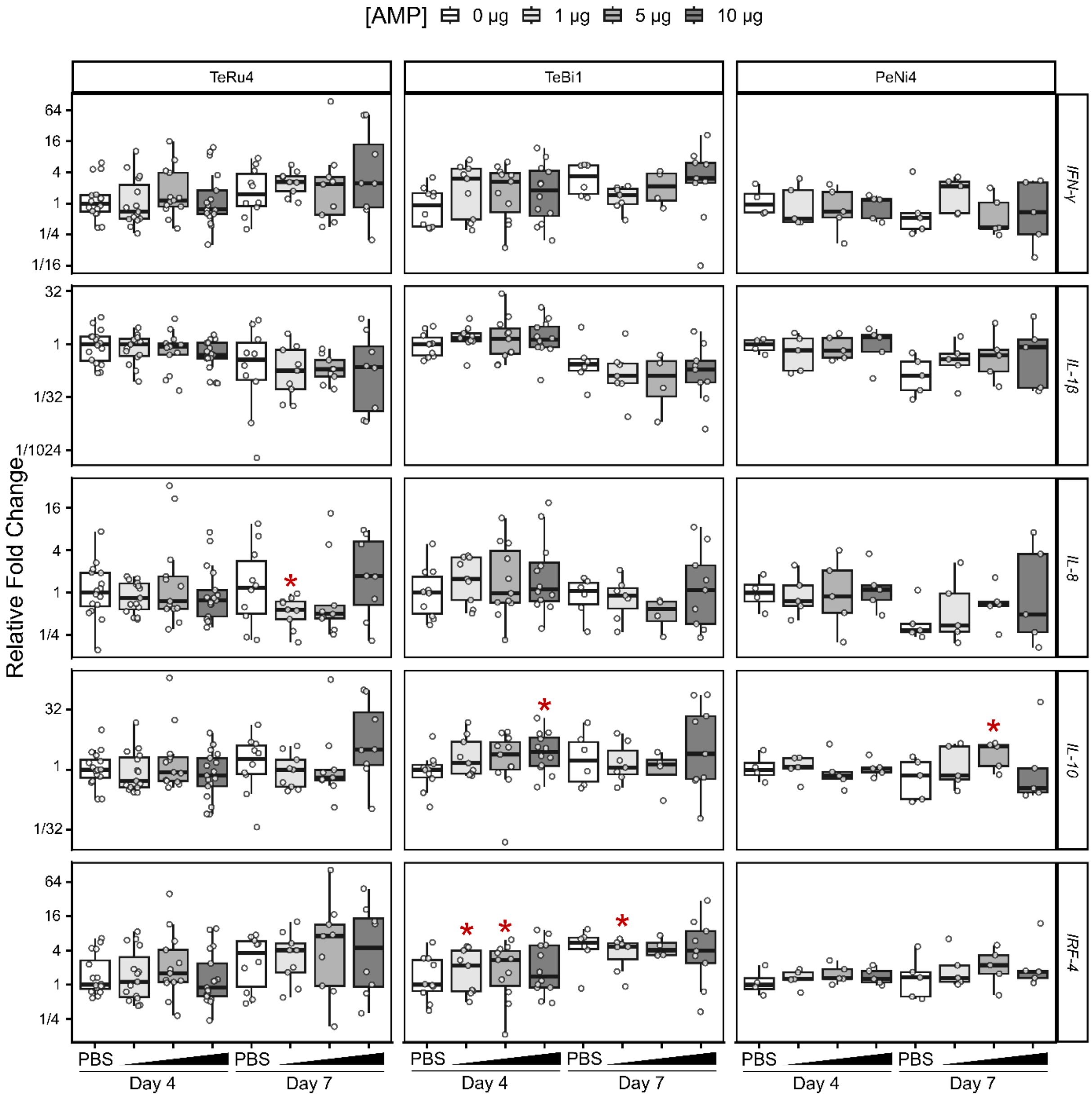


**Supplementary Figure S6.** Relative fold change values of cytokine transcripts in the spleen of avian pathogenic *E. coli* (APEC) challenged birds treated before hatch with 1, 5, or 10 µg antimicrobial peptide (AMP). Further details are in the Figure 5 legend.

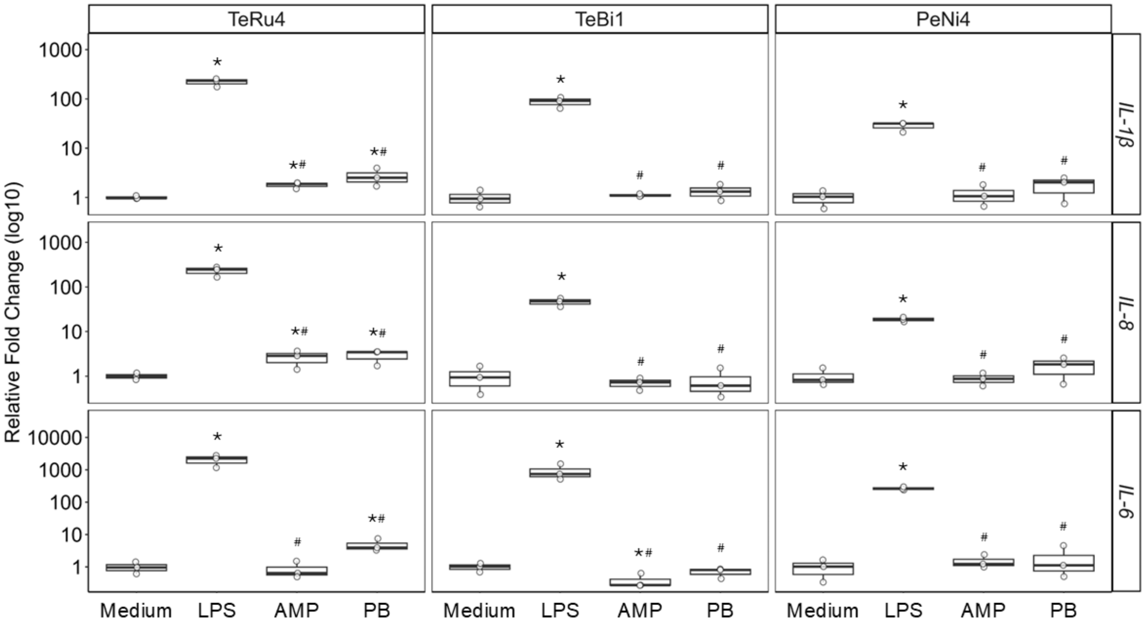


**Supplementary Figure S7.** Relative fold change of *Interleukin (IL)-1β*, *IL-8*, and *IL-6* in HD11 cells incubated for 6 h with medium, 25 ng/mL lipopolysaccharide (LPS), 16 µg/mL antimicrobial peptide (AMP; TeRu4, TeBi1, or PeNi4), or 8 µg/mL polymyxin B sulfate (PB) as determined by qPCR. A significant difference from the medium control is indicated by "*", and "#" indicates a significant difference from LPS treatment (p-value < 0.1, Mann-Whitney-U test). Open circles represent the fold change of one technical replicate relative to the medium control. The horizontal lines represent the median of the three technical replicates within each treatment condition, the boxes show the interquartile range, and the vertical whiskers represent the minimum and maximum values. Shown are representative experiments of three biological replicates.

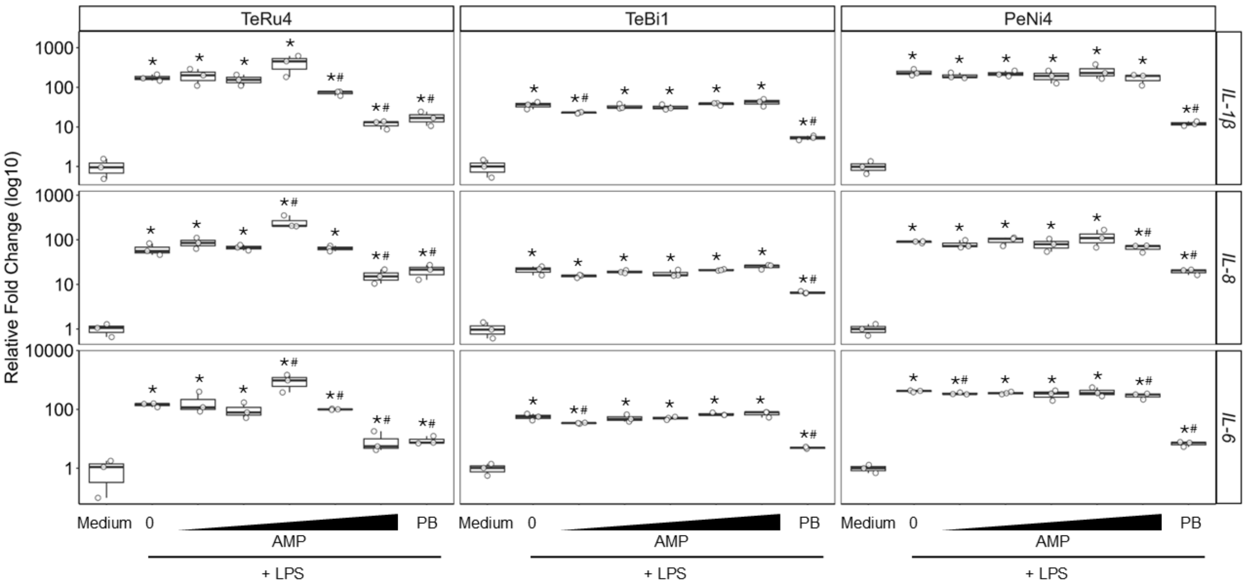


**Supplementary Figure S8.** Relative fold change of *Interleukin (IL)-1β*, *IL-8*, and *IL-6* from HD11 cells as determined by qPCR. Cells were incubated for 3 h with medium, a range of 2-fold serially diluted antimicrobial peptide (AMP; 1 to 16 µg/mL TeRu4, TeBi1, or PeNi4), or 8 µg/mL polymyxin B sulfate (PB), followed by the addition of 25 ng/mL lipopolysaccharide (LPS) for an additional 3 h. Additional details are available in the Supplementary Figure S7 legend.

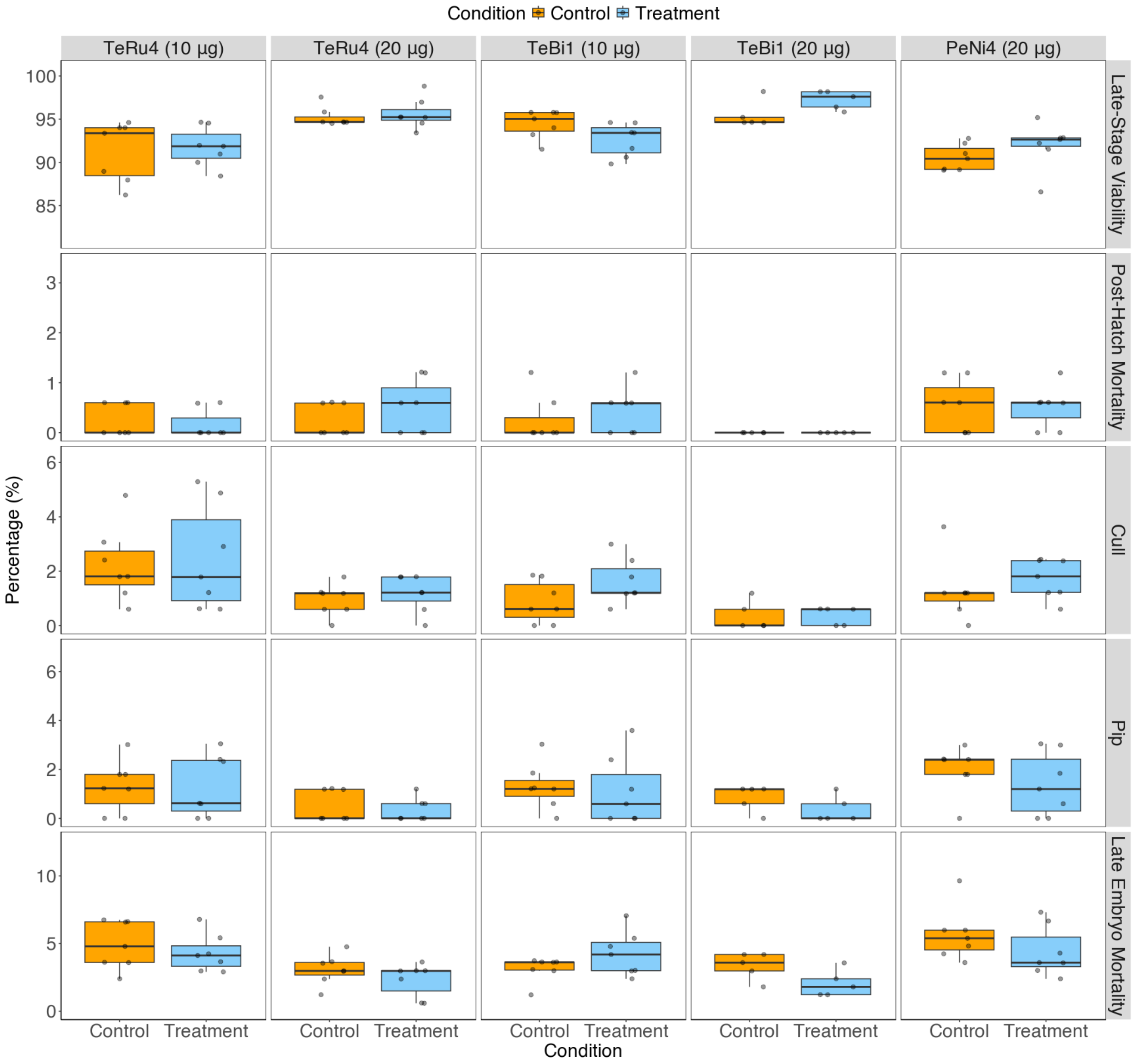


**Supplementary Figure S9.** Raw percentage values for hatchability parameters across various antimicrobial peptides (AMP) and dosages. Columns represent AMPs at specific dosages, and rows represent individual hatchability parameters. Each AMP–dosage combination corresponds to a separate pen trial. The control (orange) and AMP treatment (blue) groups are shown. Each point corresponds to a tray of eggs (n = 7 per group; n = 5 for 20 μg TeBi1). The horizontal lines represent the median of the three technical replicates within each treatment condition, the boxes show the interquartile range, and the vertical whiskers represent the minimum and maximum values. No significant differences were detected using Wilcoxon rank-sum tests.

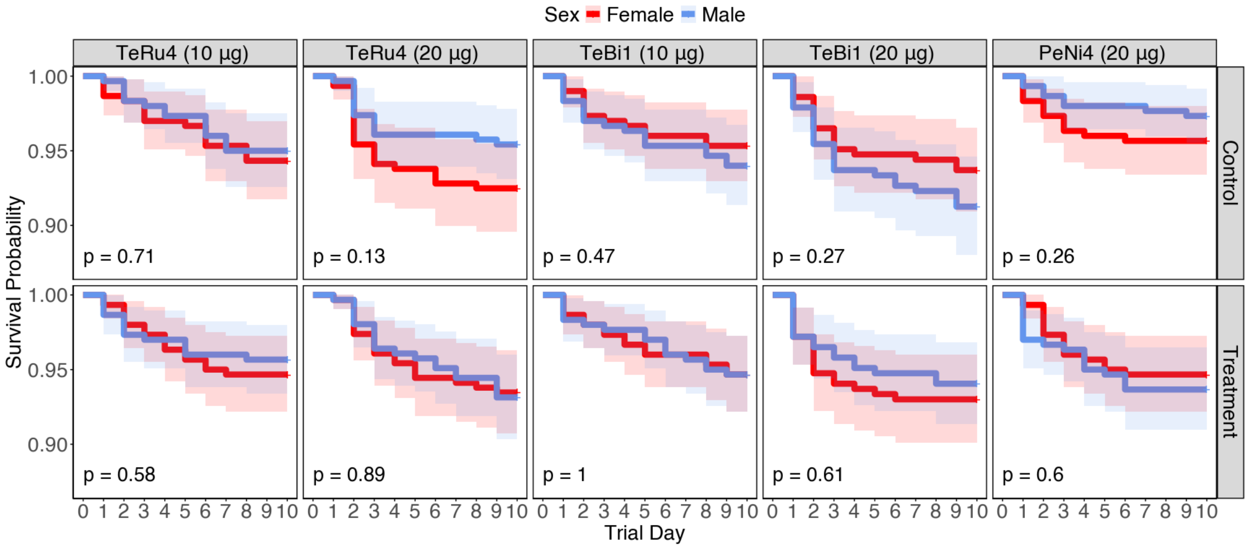


**Supplementary Figure S10.** Kaplan-Meier curves depicting the survival probabilities of the bird sexes during the first 10 days of the pen trials across different antimicrobial peptides (AMP) and dosages with sexes separated (female, red line; male, blue line). Columns represent AMPs at specific indicated dosages, and rows distinguish between control and treatment groups. Each panel represents a separate pen trial. P-values, as determined by log-rank tests, are displayed in the bottom-left corner of each panel.

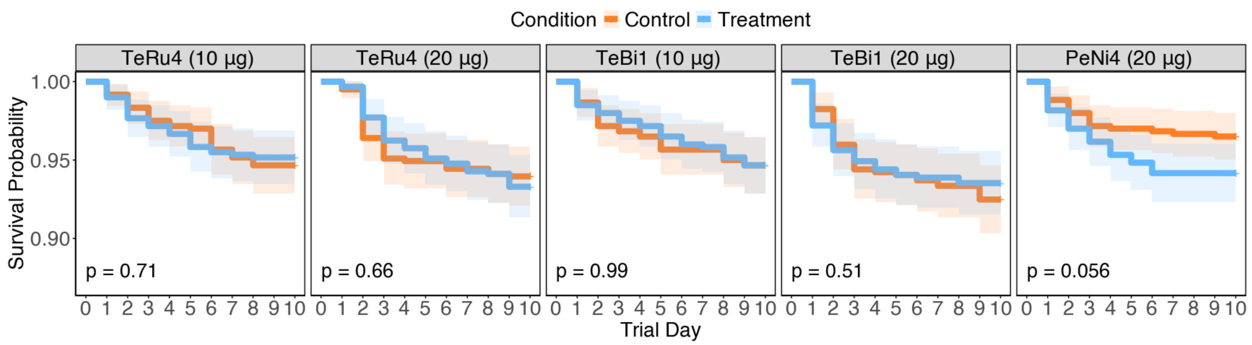


**Supplementary Figure S11.** Kaplan-Meier curves depicting the survival probabilities of control and treatment groups during the first 10 days of the pen trials across different antimicrobial peptides (AMP) and dosages with sexes combined as sexes were not significantly different from one another. The control (orange) and AMP treatment groups (blue) are shown. See Supplementary Figure S7 for details.

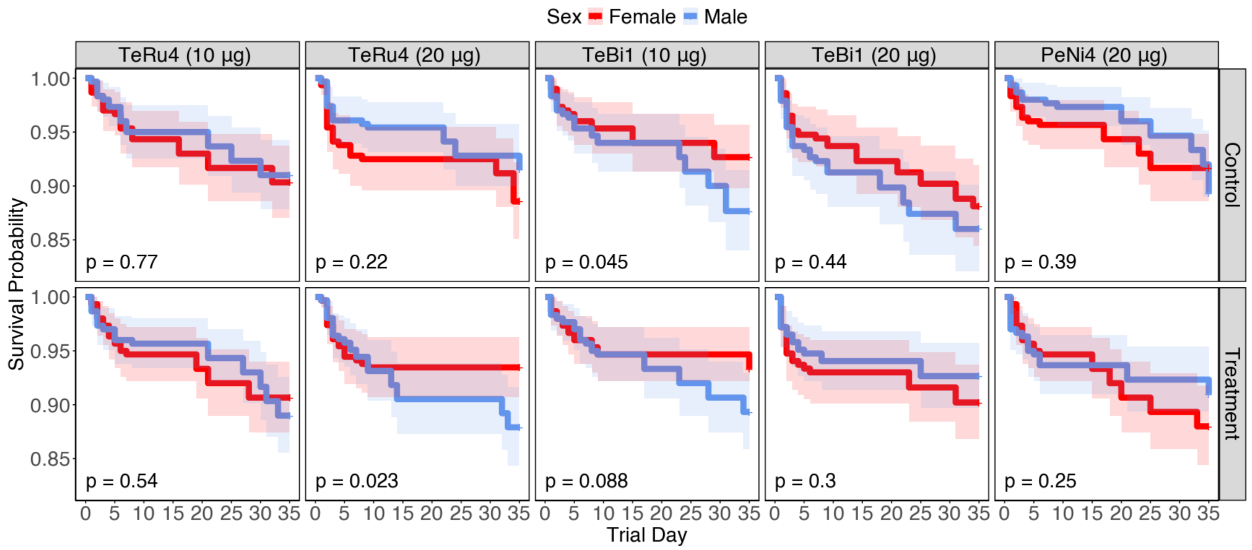


**Supplemental Figure S12.** Kaplan-Meier curves depicting the survival probabilities of female verses male chickens within a treatment condition over the full 35-day duration of pen trials across different antimicrobial peptides (AMP) and dosages. See Supplementary Figure S6 for details.

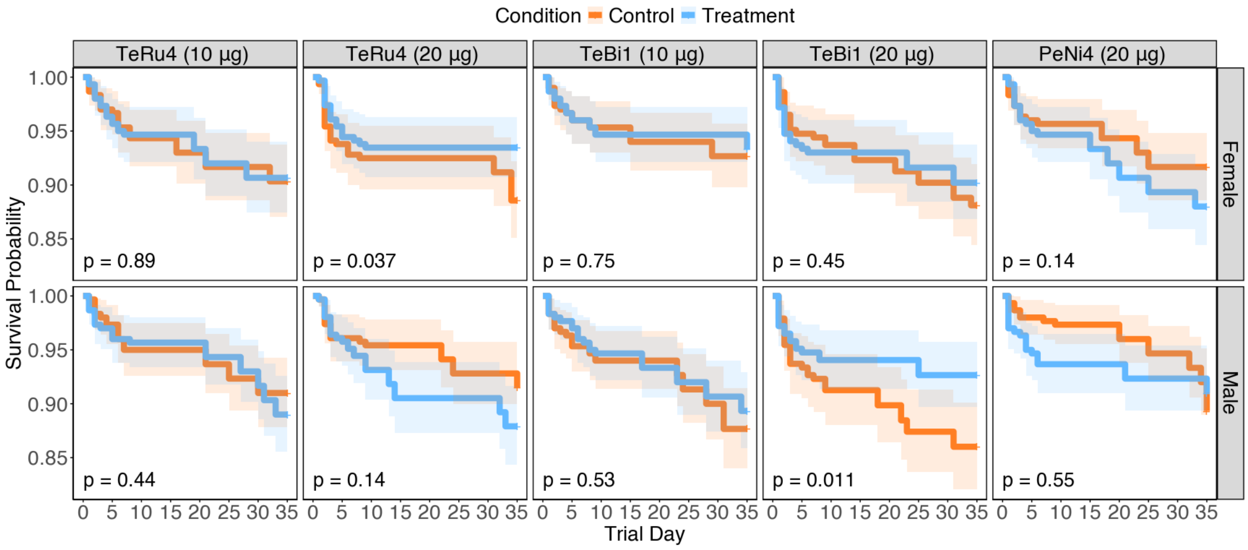


**Supplementary Figure S13.** Kaplan-Meier curves depicting the survival probabilities of experimental groups by sex over the full 35-day duration of pen trials across different antimicrobial peptides (AMP) and dosages. The control (orange) and AMP treatment groups (blue) are shown. See Supplementary Figure S7 for details.

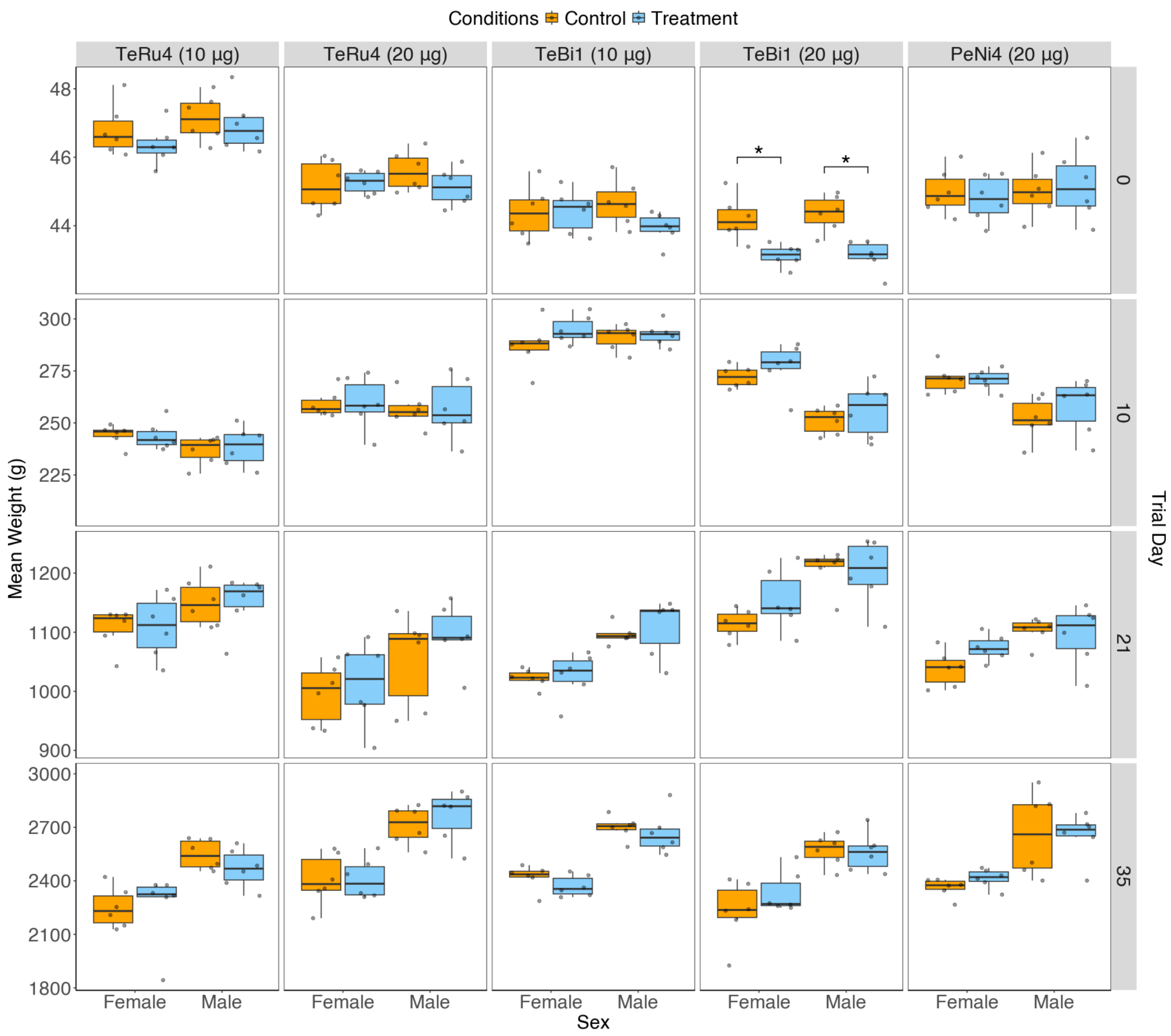


**Supplementary Figure S14.** Distribution of mean weights of female and male birds during pen trials across different antimicrobial peptides (AMP) and dosages. The control (orange) and AMP treatment groups (blue) are shown. Each point represents the mean weight for a single mini-pen (n = 6 mini-pens per group). Significant differences ($p<0.05$) between control and treatment groups within a sex, as determined by Wilcoxon rank-sum tests, are indicated by an asterisk. Additional details can be found in the Supplementary Figure S5 legend.

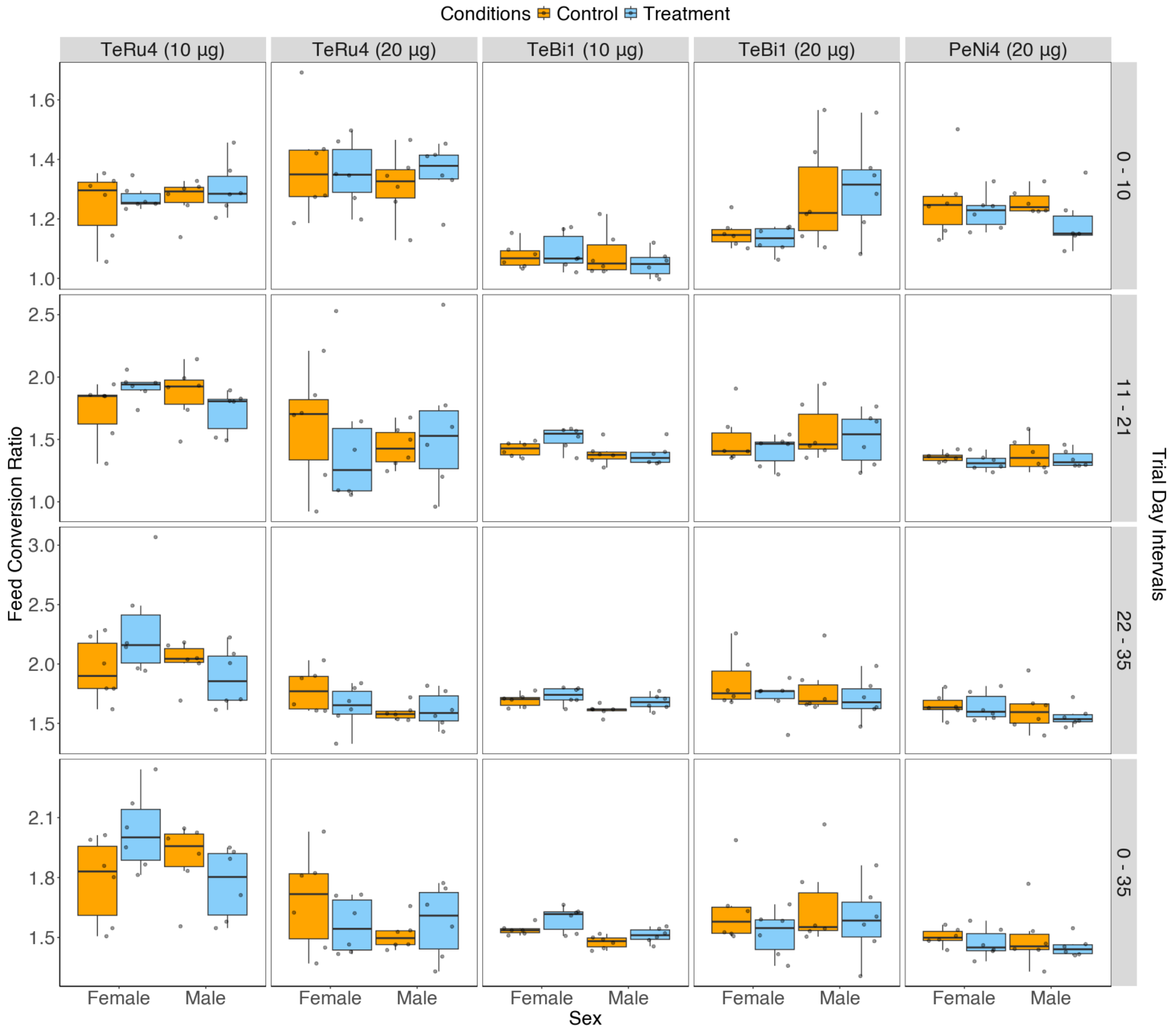


**Supplementary Figure S15.** Feed conversion ratio (FCR) of male and female birds during pen trials across different antimicrobial peptides (AMP) and dosages. The control (orange) and AMP treatment groups (blue) are shown. Each point represents the FCR for a single mini-pen (n = 6 mini-pens per group). No significant differences were detected using Wilcoxon rank-sum tests. Additional details can be found in the Supplementary Figure S5 legend.

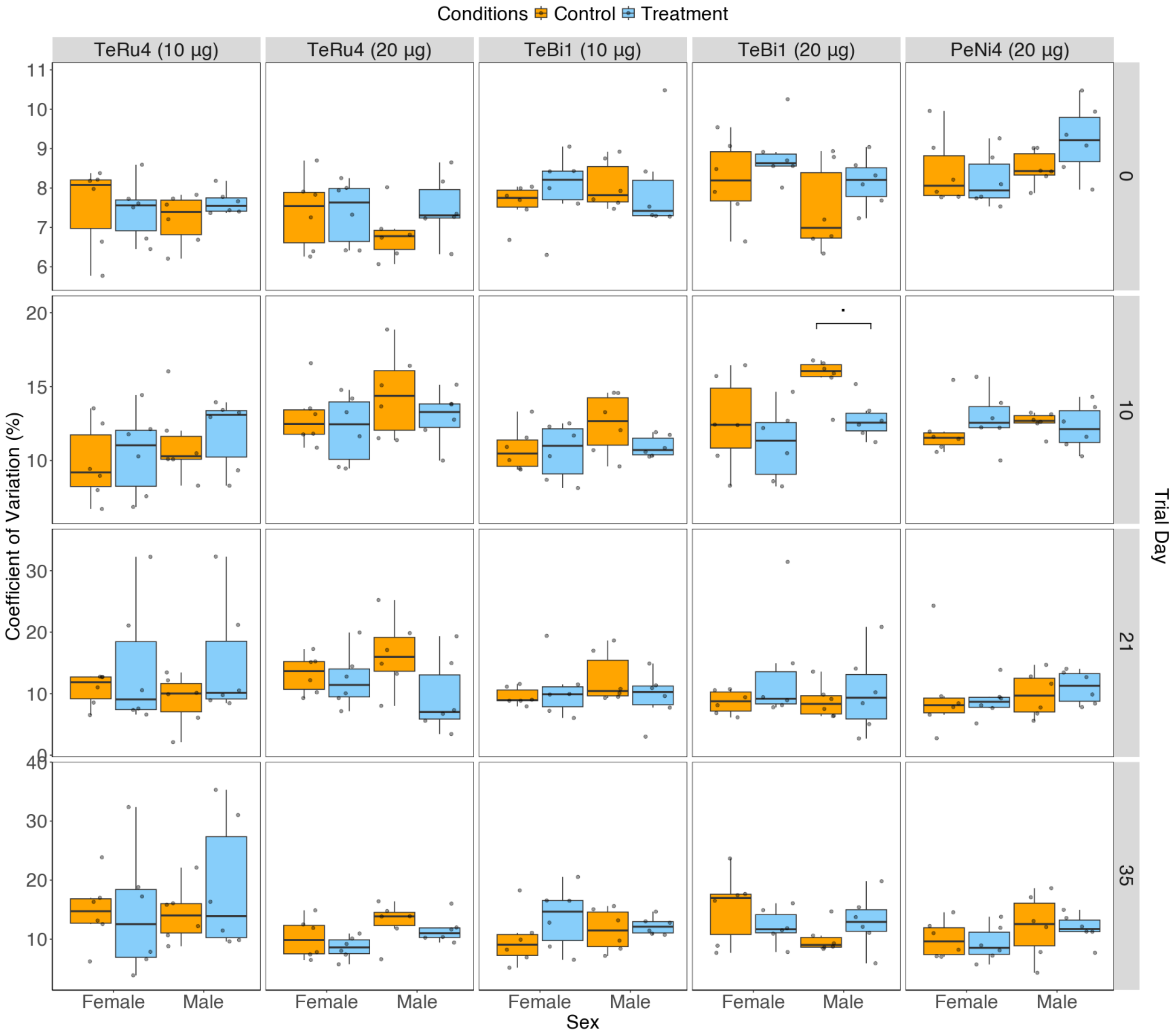


**Supplementary Figure S16.** Flock uniformity of male and female birds during pen trials across different antimicrobial peptides (AMP) and dosages. The control (orange) and AMP treatment groups (blue) are shown. Each point represents the % CV for a single mini-pen (n = 6 mini-pens per group). Trends ($0.05<p<0.1$) between control and treatment groups within a sex, as determined by Wilcoxon rank-sum tests, are indicated by a dot. Additional details can be found in the Supplementary Figure S5 legend.

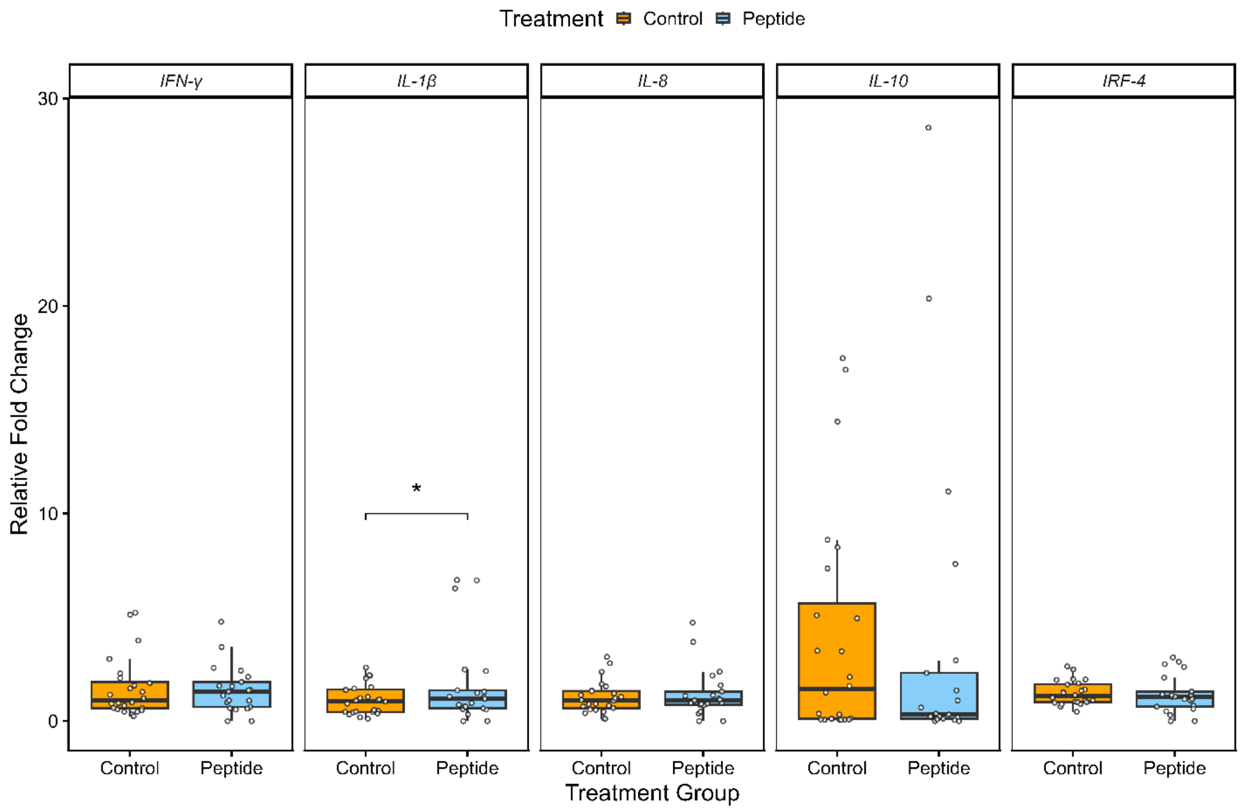


**Supplementary Figure S17.** Relative fold change values of cytokine transcripts in the spleens of pen trial birds (no pathogen challenge) treated with 10 µg TeBi1. Additional details in the figure 6 legend.
